# Structural and functional insights into the type III-E CRISPR-Cas immunity

**DOI:** 10.1101/2022.08.22.504715

**Authors:** Xi Liu, Laixing Zhang, Hao Wang, Yu Xiu, Ling Huang, Zhengyu Gao, Ningning Li, Feixue Li, Weijia Xiong, Teng Gao, Yi Zhang, Maojun Yang, Yue Feng

**Author notes:** These authors contributed equally to this work (Xi Liu, Laixing Zhang, Hao Wang, Yu Xiu, Ling Huang, Zhengyu Gao). Correspondence (L. Zhang), (M. Yang), (Y. Feng).

## Abstract

The type III-E CRISPR-Cas system comprises a Cas effector (gRAMP), a TPR-CHAT and several ancillary proteins. However, both the structural features of gRAMP and the immunity mechanism remain unknown for this system. Here, we report a series of structures of gRAMP-crRNA, either its alone or in complex with target RNA or TPR-CHAT (called Craspase), and Craspase complexed with cognate (CTR) or non-cognate target RNA (NTR). Importantly, the 3’ anti-tag region of NTR and CTR bind at two distinct channels in the Craspase, and CTR with a non-complementary 3’ anti-tag induces a marked conformational change of the TPR-CHAT, which allosterically activates its protease activity to cleave an ancillary protein Csx30. This cleavage then triggers an abortive infection as the antiviral strategy of the type III-E system. Together, our study provides crucial insights into both the catalytic mechanism of the gRAMP and the immunity mechanism of the type III-E CRISPR-Cas system.

---

In prokaryotes, CRISPR-Cas (clustered regularly interspaced short palindromic repeats and its associated genes) adaptive immune systems fight against invading nucleic acids from bacteriophages and plasmids (1–3). CRISPR-Cas systems are divided into class 1 and 2 based on the architecture of the Cas effector. Class 1 systems (further divided into types I, III, and IV) are represented by their multi-subunit effector complexes, while class 2 systems (type II, V, and VI) are composed of single-subunit protein effectors. Out of these types, type III CRISPR-Cas systems are of particular interest due to their delicate mechanisms and cleavage of both RNA and DNA of the invaders (4,5). Type III system recognizes invading RNA molecules as target and is further divided into subtypes III-A to III-F (6). Type III system is characterized by the signature Cas10 protein in the locus (7), which comprises an N-terminal HD nuclease domain, two Palm domains and other domains. Type III-A/D systems encode the Csm complex, composed of Csm1-5 and crRNA. Type III-B/C systems encode the Cmr effector complex, which consists of Cmr1-6 and crRNA. The Cas10 subunit is essential for the canonical type III systems, in which recognition of an RNA target with mismatching protospacer flanking sequence (PFS) activates the non-specific single-stranded DNA cleavage by the Cas10 HD domain, and cyclic oligoadenylate (cOA) generation by the Cas10 Palm domains (4,8–10). cOAs could further act as second messengers to activate CRISPR-Cas associated proteins such as the Csm6/Csx1 family RNases (8,10–14). Activation of both activities of the Cas10 subunit requires the 5’ repeat region (named 5’ tag) of crRNA and 3’ sequence flanking the target RNA sequence (named 3’ anti-tag) to be non-complementary (called cognate target RNA, CTR). In contrary, both activities will be repressed if these flanking sequences are complementary (non-cognate target RNA, NTR), possibly to avoid autoimmune response upon recognition of self-transcripts (8–10,15).

However, the type III-E system is largely different from the typical type III-A to III-D systems. On the one hand, in the type III-E system, four Cas7 domains and a Csm2/Cmr5-like small subunit (Cas11) are fused into one single large protein, named gRAMP (6), like the large multi-domain proteins of class 2 CRISPR-Cas systems. On the other hand, the *cas10* and *csm4*/*cmr5* genes are absent from the CRISPR locus of type III-E system, suggesting its different immune mechanisms. Very recently, two studies uncovered that the effector of the type III-E system is able to process pre-crRNA into mature crRNA and cleave target RNA at two defined positions six nucleotides apart (16,17). However, unlike the Cas13 family, which also processes pre-crRNA and targets RNA molecules (18,19), the type III-E effector does not show collateral cleavage activity towards non-specific RNA molecules. Meanwhile, the III-E effector was also engineered to mediate RNA knockdown and editing in mammalian cells, with no effects on cell viability (17). Therefore, the type III-E effector represents a good candidate for new programmable RNA-targeting tools free of collateral activity and cell toxicity. Interestingly, the type III-E effector was found to stably bind a caspase-like TPR (tetratricopeptide repeat)-CHAT (Caspase HetF Associated with TPRS) peptidase to form a larger complex, named “Craspase” (CRISPR-guided caspase), which has been suggested to work as an activated protease induced by recognition of target RNA by the type III-E effector (16). However, no such activity has been verified and no substrate has been identified for the TPR-CHAT peptidase. Moreover, the mechanisms underlying the other features of the type III-E gRAMP have also remained enigmatic.

## Overall structure of the gRAMP-crRNA complex

To understand the molecular basis of how the architecture of gRAMP determines its function, we purified recombinant gRAMP-crRNA from *Candidatus* “Scalindua brodae” (*Sb*-gRAMP-crRNA, short for gRAMP-crRNA hereafter), by co-expressing the gene with a plasmid containing five copies of the first spacer-repeat from the native CRISPR array. Then we solved the cryo-EM structure of gRAMP-crRNA at 3.01 Å resolution (Figure 1A and 1B and Fig. S1; Table S1). Previous bioinformatic analysis indicated that type III-E effectors comprise one Cas11-like domain, and four Cas7-like domains, the last one with a large insertion (16,17). In our structure, gRAMP contains the above mentioned domains and one crRNA molecule, except that the insertion domain within the last Cas7-like domain displays a poor density (Fig. S1). While gRAMP maintains the overall shape of type III CRISPR-Cas effectors (20–28), it exhibits several unique features compared to Csm and Cmr complexes except the absence of the signature Cas10 subunit. Notably, gRAMP does not display the typical conformation of central double-helical core in type III-A/B complexes. In gRAMP, four Cas7-like domains (named as Cas7.1-Cas7.4, hereafter) packs with each other in their sequence order in the protein, wrapping the crRNA from its 5’ to 3’ end in the order of Cas7.1 to Cas7.4 (Figure 1B). The Cas11-like domain folds majorly as a five-α-helical bundle with a short α3 helix (Figure 1C), and packs against Cas7.2 and Cas7.3 domains. Out of the four Cas7 domains, the Cas7.1-7.3 domains are overall structurally more similar to each other, and the Cas7 family protein Csm3 in the type III-A interference complex is their most similar structural homolog (Figure 1C and Fig. S2, A-D). Cas7.1-7.3 domains exhibit typical features of Cas7-like domains, with fingers-, palm-, web- and thumb-shaped sub-domains, and Cas7.2 and Cas7.3 are structurally more similar to each other (Figure 1C and Fig. S2). While Dali search also returned Csm3 as the most similar structure for Cas7.4 domain, it also displays features not conserved in Csm3 homologs, such as striking structural variations in the typical “β-thumb” sub-domain, multiple extra motifs as well as incorporation of the insertion domain (Figure 1C and Fig. S2F). Each of the Cas7 domains coordinates a zinc ion with a zinc finger motif (Fig. S2G). The small Cas11-like domain (residues 266-375) is linked to Cas7.1 and Cas7.2 through two long loops (Fig. S3). The crRNA passes through the entire gRAMP effector, suggesting its essential role in sustaining the conformation and stability of gRAMP. This explains why expressing *Sb*-gRAMP in the absence of pre-crRNA yields a lower amount and more degraded forms of the protein, as shown in the previous study (16). While the repeat and spacer in the CRISPR array are designed 36 and 38 base pairs, respectively, the gRAMP only contains 18-nt traceable repeat region (5’ tag) and 17-nt traceable spacer region (Fig. S2E). Consistently, previous study also showed that the sequences of the last 14-nt of the crRNA repeat are conserved (16). The spacer region of the crRNA is bound by the interlocking Cas7.2-7.4 domains, and locked at two sites by the “thumbs” of Cas7.2 and 7.3, respectively (Figure 1D). This binding mode results in the typical “5+1” pattern, i.e., 5 consecutive bases are followed by a 1 base gap, showing two “kinks” at A4 and U10 of the crRNA, respectively (Figure 1D).

**Figure 1.**
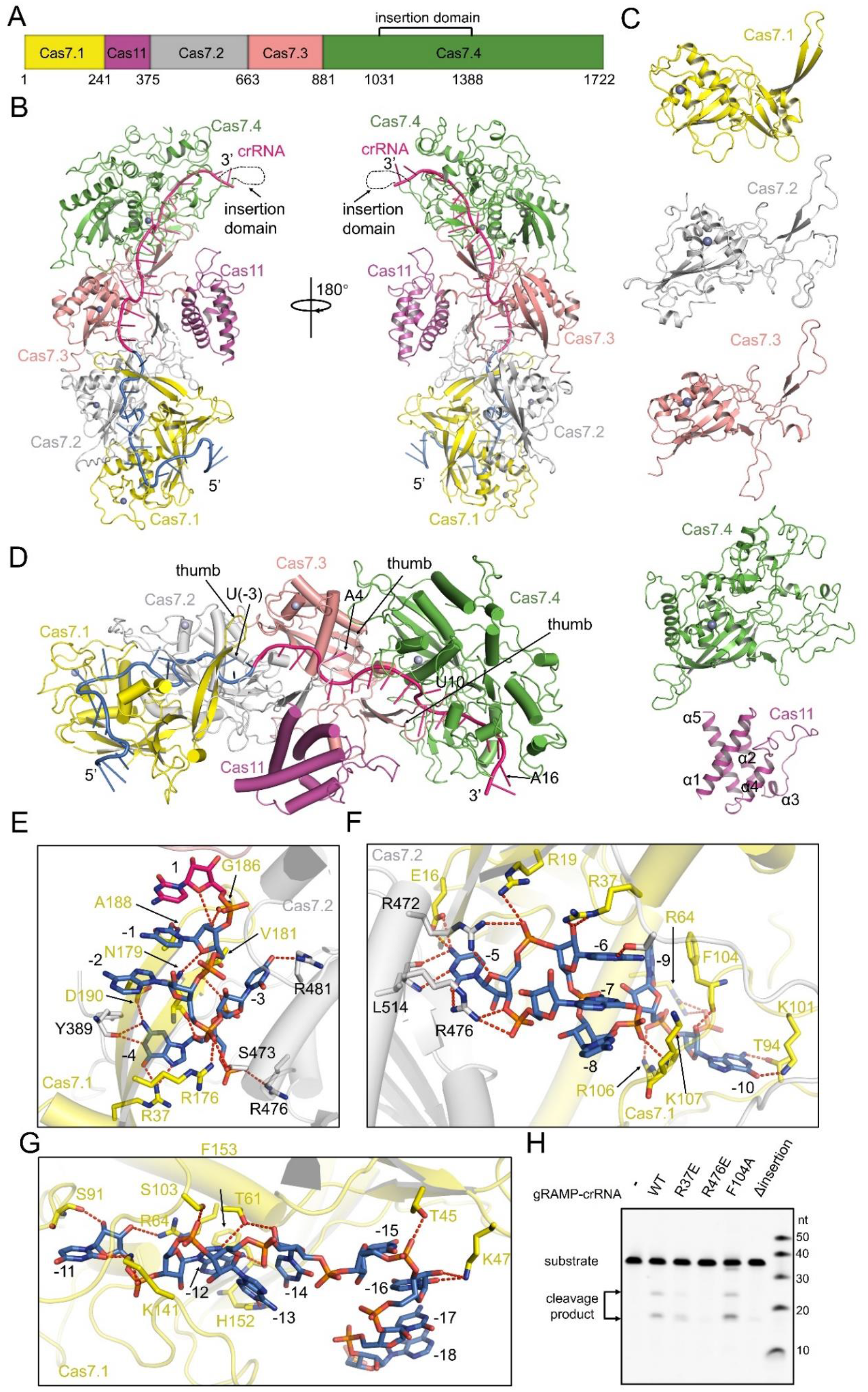
Cryo-EM structure of the gRAMP-crRNA. (A) Domain architecture of the gRAMP in *Candidatus* “Scalindua brodae”. The Cas7.1-7.4 and Cas11 domains are colored in magenta, yellow, gray, pink and green, respectively. (B) Overall structure of the gRAMP-crRNA. The domains of gRAMP are colored as in (A). The crRNA is colored in hot pink and marine, for the spacer and repeat region, respectively. The unmodelled insertion domain within Cas7.4 is marked. (C) Structures of separate domains of the gRAMP. Secondary structures are labeled for the Cas11 domain. (D) Close-up view of the binding of the crRNA. The thumbs of the Cas7.1-7.3 domains and the kinked nucleotides in the crRNA are marked. (E-G) Interaction between Cas7.1/7.2 domains of the gRAMP and the nucleotides (−1)-(−4) of crRNA (E), (−5)-(−10) of crRNA (F) or (−11)-(−18) of crRNA, respectively. Red dashed lines represent polar interactions. (H) The target RNA cleavage with gRAMP-crRNA and its mutants. ΔInsertion indicates gRAMP-crRNA with the insertion domain (residues 1031-1388) deleted and substituted by a GSG linker.

## The insertion domain is indispensable for the RNase activity of gRAMP

The 18-nt 5’ tag including the conserved U(−14)-C(−1) of crRNA is held in the Cas7.1-7.2 domains (Figure 1D). Detailed interactions between the 5’ tag of crRNA and Cas7.1/7.2 domains are shown in Figure 1E-G and Fig. S4A. Biochemical studies showed that R37E and R476E (Figure 1E and 1F) mutations in gRAMP reduced crRNA-dependent RNA cleavage, suggesting the role of 5’ tag binding in the nuclease activity of gRAMP (Figure 1H). The 17-nt traceable spacer region is held tightly in the channel formed in the Cas7.2-7.4 domains mainly through interactions involving its phosphate/ribose backbones (Figure 1B and Fig. S4A), consistent with the variable sequences among crRNA spacers (16,17). The detailed interactions between the spacer region of crRNA and gRAMP are shown in Fig. S4A. Notably, while the region of insertion domain of Cas7.4 displays a poor density, its location indicates that it interacts with the 3’ end of traceable region of crRNA and is colocalized with the rest of the spacer region (Figure 1B and Fig. S1D). To investigate the role of this insertion domain, we co-expressed the CRISPR array and a gRAMP mutant with the insertion domain deleted (gRAMPΔinsertion) and purified it similarly as WT protein. However, the ratio of 254/280 nm absorbance of the purified gRAMPΔinsertion is much lower than that of WT gRAMP (Fig. S5A), and no RNA was detected in the purified gRAMPΔinsertion protein (Fig. S5B). Consistently, target RNA cleavage assay showed that gRAMPΔinsertion displays almost no cleavage activity (Figure 1H).

Interestingly, during the preparation of our study, a recent study into *Di*-gRAMP (from *Desulfonema ishimotonii*) indicated that the insertion domain is dispensable for the catalytic activity of *Di*-gRAMP (29) and their study identified H43 of Cas7.1 domain is responsible for pre-crRNA processing between (−15) and (−16) of the repeat region. However, this residue is not conserved in *Sb*-gRAMP (Fig. S3), suggesting a different processing mechanism. Notably, the previous study reported that when co-expressing with CRISPR arrays, *Sb*-gRAMP contains a crRNA with 27-28 nt repeat and 16-25 nt spacer (16), both of which are shorter than their exact sequence, suggesting that processing may happen at both the repeat and spacer region. Pre-crRNA processing experiment showed that the gRAMPΔinsertion displays a weakened processing activity compared to WT protein (Fig. S5C), suggesting that the insertion domain is involved in the pre-crRNA processing of *Sb*-gRAMP, at least in the spacer region based on the location of this domain. The differences in the function of the insertion domain and pre-crRNA processing between *Sb*-gRAMP and *Di*-gRAMP may reflect the mechanistic differences between type III-E effectors from different species.

## Overall structure of the gRAMP-crRNA complexed with target RNA

To investigate how the gRAMP-crRNA recognizes and cleaves target RNA, we determined the cryo-EM structure of gRAMP-crRNA bound to target RNA (TR) complementary to the spacer region (38-nt) at 2.89 Å resolution (Figure 2A and 2B; Fig. S1; and Table S1). The spacer-target RNA duplex displays a discontinuous dimer structure. Structural alignment between gRAMP-crRNA-TR and the apo gRAMP-crRNA reveals overall similarity (root mean square deviation, RMSD value of 0.35 Å among 992 Cα atoms) but a notable rigid-body rotation of the Cas11 domain (Figure 2C-E). Here, we focus on how target RNA is recognized and cleaved. As in type III-A/B complexes, the Cas11 domain offers a positively charged surface to bind TR (Figure 2F), which base pairs with crRNA to form a duplex with positions corresponding to A4 and U10 in crRNA flipped out (Fig. S4B). Stacking interactions and polar interactions between the backbone phosphate/ribose groups of the two RNA molecules and gRAMP are involved in the stabilization of the RNA duplex (Fig. S4B). The flipped-out bases U4’ and A10’ are stabilized by R294 and R323 from the Cas11 domain, respectively (Figure 2G). Mutational studies showed that disruption of gRAMP-target-RNA binding in the spacer region (Fig. S4B) also reduced crRNA-dependent RNA cleavage (Figure 2H). Notably, the Y367A mutation designed to disrupt binding of backbone phosphate of U3’ of target RNA only affects cleavage at this specific site (Figure 2G and 2H).

**Figure 2.**
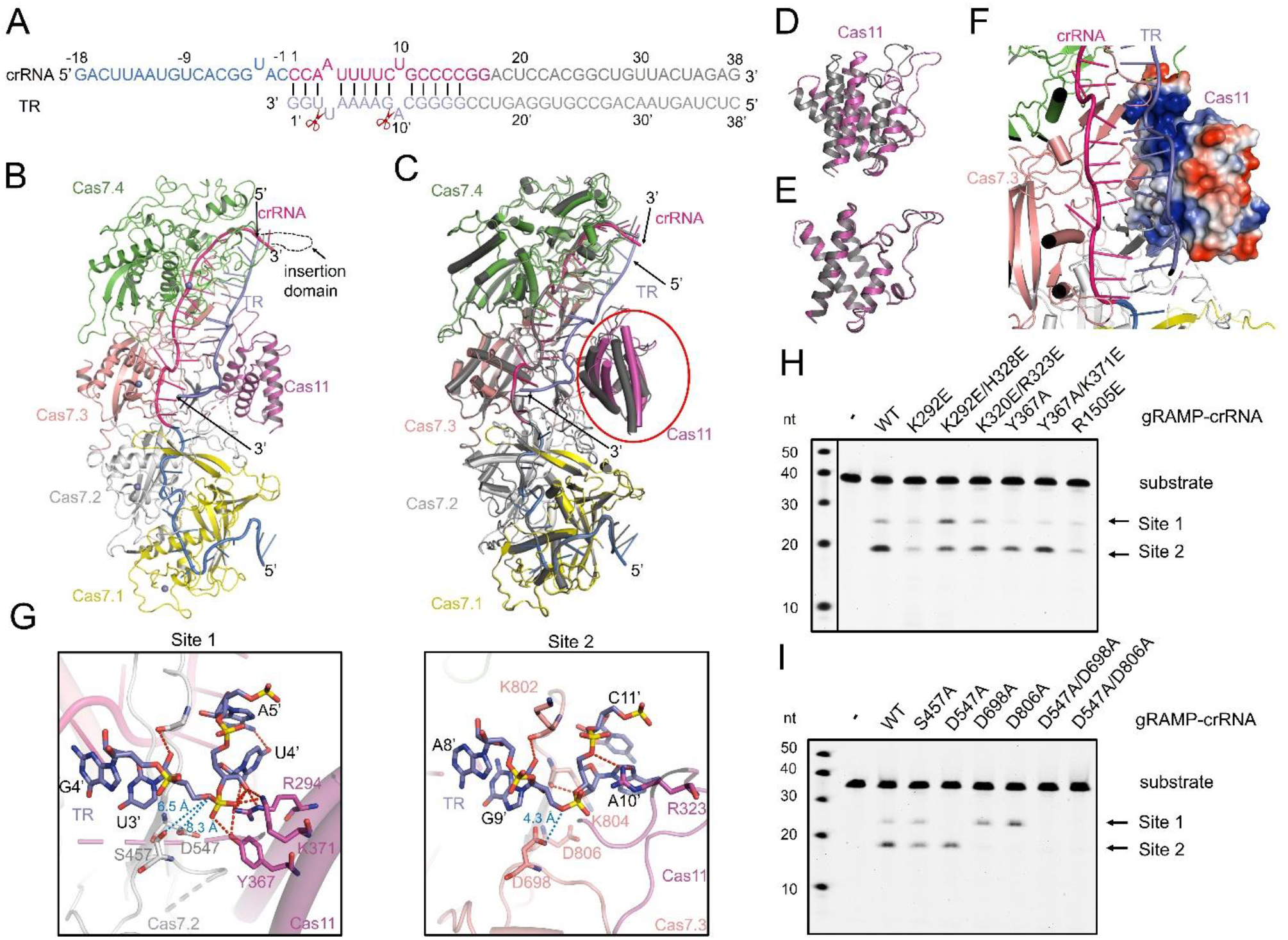
Structure of the gRAMP-crRNA-TR and RNA cleavage mechanism. (A) Schematic representation of the crRNA-TR duplex. The nucleotides of crRNA and TR used in the experiment are depicted in lines. The TR is colored in slate. The modelled nucleotides of crRNA and TR are shown as in the structure. The invisible nucleotides are colored in gray. The scissors indicate cleavage sites. (B) Overall structure of the gRAMP-crRNA-TR. TR is colored as in (A). (C) Structural superimposition of the gRAMP-crRNA (colored dark gray) and gRAMP-crRNA-TR (colored as in B). The region of Cas11 domain is highlighted in a circle. (D) Close-up view of the region of Cas11 domain within the structural alignment in (C). (E) Structural alignment of the separate Cas11 domain in gRAMP-crRNA and gRAMP-crRNA-TR. (F) Close-up view of the Cas11 region in the gRAMP-crRNA-TR structure, in which the Cas11 domain is colored in electrostatic model. (G) Detailed interactions between gRAMP and TR in the two cleavage sites. Red dashed lines represent polar interactions. (H) The target RNA cleavage with gRAMP-crRNA and its mutants impairing TR binding in the spacer region. The product bands representing cleavage at Site1 and 2 are marked. (I) The target RNA cleavage with gRAMP-crRNA and its mutants with mutations at two cleavage active sites.

## Target RNA cleavage mechanism of gRAMP-crRNA

Next, we investigated how gRAMP-crRNA cleaves target RNA at positions U3’-U4’ (Site 1) and G9’-A10’ (Site 2) of TR (16). Previous study reported that D698 of *Sb*-gRAMP is responsible for the cleavage at Site 2. Consistently, in the gRAMP-crRNA-TR structure, D698 in the catalytic loop from the Cas7.3 domain is located right under the scissile bond, with a distance of ~4.3 Å as measured from the sidechain carboxyl group oxygen of D698 to the scissile P-O3’ phosphate oxygen at Site 2 (Figure 2G, right panel). However, the catalytic residue for Site 1 has not been identified due to the low sequence identity between gRAMP and Csm3/Cmr4. Interestingly, detailed investigation of the gRAMP residues with polar sidechains near Site 1 showed that, S457 in the catalytic loop of Cas7.2 domain is at the corresponding position as D698 of Cas7.3 domain with a distance of ~6.5 Å as measured from the sidechain hydroxyl group oxygen of S457 to the scissile P-O3’ phosphate oxygen (Figure 2G, left panel). However, the previous study showed that mutation of S457 does not affect RNA cleavage (16), which was also confirmed by our study (Figure 2I). Notably, D547 in the “thumb” of Cas7.2 domain is also near the scissile bond, with a distance of ~8.3 Å as measured from the sidechain carboxyl group oxygen of D547 to the same scissile phosphate oxygen (Figure 2G, left panel). Further RNA cleavage assays showed that the D547A mutation abolishes cleavage at Site 1, and the D547A/D698A mutant exhibits no cleavage at either site (Figure 2I). Inspired by this, we also tested the D547 corresponding site in the Cas7.3 domain, D806, whose mutation showed the same phenotype as the D698 mutation in RNA cleavage assay (Figure 2I). Taken together, our results suggested that D547 is responsible for cleavage at Site 1, and D698/D806 are responsible for cleavage at Site 2 in *Sb*-gRAMP.

## Structure of the Craspase complex

To investigate the architecture and mechanism of the Craspase complex, we purified the gRAMP-crRNA complexed with TPR-CHAT to homogeneity, and solved its cryo-EM structure at 2.88 Å resolution (Figure 3A; Fig. S1; and Table S1). The structure of gRAMP-crRNA in the Craspase remains largely unchanged compared with its apo form with an RMSD of 0.42 Å among 1279 Cα atoms (Fig. S6A). Notably, two loops (G376-D386 and S449-N453) of gRAMP, whose densities are poor in its apo structure, become well ordered in the Craspase structure, possibly resulting from their involvement in the gRAMP-TPR-CHAT interaction (Fig. S6B). In the Craspase, TPR-CHAT interacts exclusively with the gRAMP but not the crRNA, with a buried surface area of ~3435.1 Å^2^. The TPR-CHAT contains an N-terminal TPR domain (residues 1-363), and a C-terminal CHAT domain (residues 388-716) (Figure 3B and Fig. S7). The density for residues linking the two domains is invisible, possibly due to its flexibility nature. Notably, in the Craspase, TPR-CHAT is bound at the region of gRAMP which holds the 5’ tag of crRNA, similar to the position of Csm1/Cmr2 in the type III-A/B CRISPR-Cas complex (Fig. S6C). TPR-CHAT mainly engages two separate regions of the Cas7.2 domain of gRAMP using its TPR and CHAT domains, respectively, and thus further locking the 5’ tag region of the crRNA between the two proteins (Fig. S6D). In both interfaces, hydrophobic interactions are primarily involved. Specifically, in the interface involving the TPR domain, F381, I383, L384 of gRAMP, densities of which are all invisible in the apo gRAMP-crRNA structure, interacts with L49, K50, I53, V78, K92 and A96 of TPR-CHAT through hydrophobic interactions (Figure 3C). Meanwhile, K42/K43 and K333/R595 (from CHAT domain) of TPR-CHAT forms potential electrostatic interactions with D746/D749 and D184 of gRAMP, respectively. In the interface involving the CHAT domain, hydrophobic interactions are primarily observed between a helix (K434-Y450) of TPR-CHAT and a loop (L401-P408) of gRAMP (Figure 3D). Consistently, mutations of some of the TPR-CHAT residues mentioned above, which are conserved among TPR-CHAT homologs (Fig. S7), abolished or decreased gRAMP binding in the MST assay (Figure 3E).

**Figure 3.**
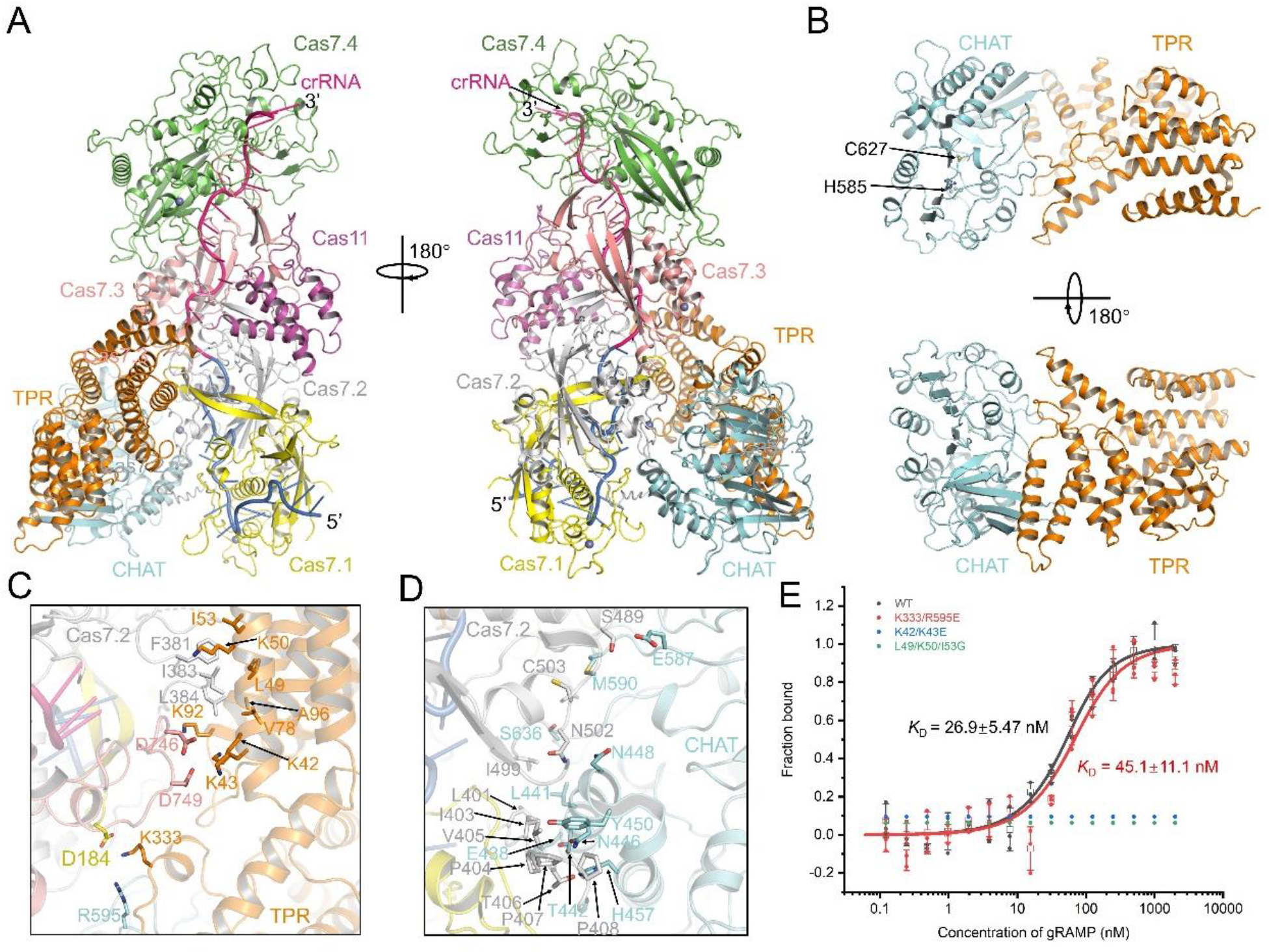
Cryo-EM structure of the Craspase. (A) Overall structure of the Craspase. The gRAMP-crRNA is colored as in Figure 1B. TPR-CHAT is colored in orange and cyan, for the TPR and CHAT domain, respectively. (B) Overall structure of TPR-CHAT in the Craspase, colored as in (A). Two active site residues are shown in sticks. (C-D) Detailed interactions between gRAMP and the TPR domain (C) and CHAT domain (D) of TPR-CHAT. (E) MST assays of the binding of gRAMP-crRNA to TPR-CHAT and its mutants. Individual values from three independent experiments are shown. Binding curves and *K*_D_ values are also shown. Error bars indicate the s.d. of three independent experiments.

## CTR binding activates the protease activity of Craspase towards Csx30

The formation of the Craspase complex suggests that binding of target RNA might activate the protease activity of TPR-CHAT in the Craspase complex, in light of the case of Csm1/Cmr2 in canonical type III CRISPR-Cas interference complexes (4). To find the potential substrate of TPR-CHAT, we first tested whether the protein products of the genes neighborhood of TPR-CHAT act as the target of TPR-CHAT. In the gene locus harboring *gRAMP* and *TPR-CHAT*, there are normally three conserved accessory genes, which are *Csx30*, *Csx31* and a sigma factor E (σ^E^) encoding gene *RpoE* (16,17). Interestingly, out of the three proteins, protease cleavage assays showed that Csx30, but not RpoE, can be cleaved by the Craspase in the presence of CTR (with non-complementary 3’ anti-tag) but not TR or NTR (with complementary 3’ anti-tag, Figure 4A and Fig. S8A). Csx31 was not tested itself because it could not be purified to homogeneity under our experimental conditions. However, Csx30 cannot be cleaved by the Craspase in which TPR-CHAT contains mutations at its active site (H585 or C627) in the presence of CTR, suggesting that the observed proteolytic activity results from the catalytic activity of TPR-CHAT (Figure 4B). SDS-PAGE analysis revealed two cleavage products with molecular weights of ~45 and ~18 kDa, respectively (Figure 4A). To investigate the cleavage site, we first confirmed that the cleavage happens near the C-terminus of Csx30 (Fig. S8B). Then, the full-length Csx30 and its two cleavage products (Csx30-N and Csx30-C) were all purified to homogeneity and subjected to mass spectrometry. The result determined that cleavage occurs after a non-polar residue L407 of Csx30 (Fig. S8C). Then we performed alanine scanning mutagenesis of the Csx30 P4-P4′ residues to determine the specificity of proteolytic cleavage of Csx30. The result showed that the L407 P1 position is essential for cleavage, and proteolysis is also inhibited by mutations that disrupt the P3, P2, P1’ and P2’ residues (Fig. S8D). Taken together, CTR binding activates the protease activity of Craspase to cleave Csx30 after its residue L407.

**Figure 4.**
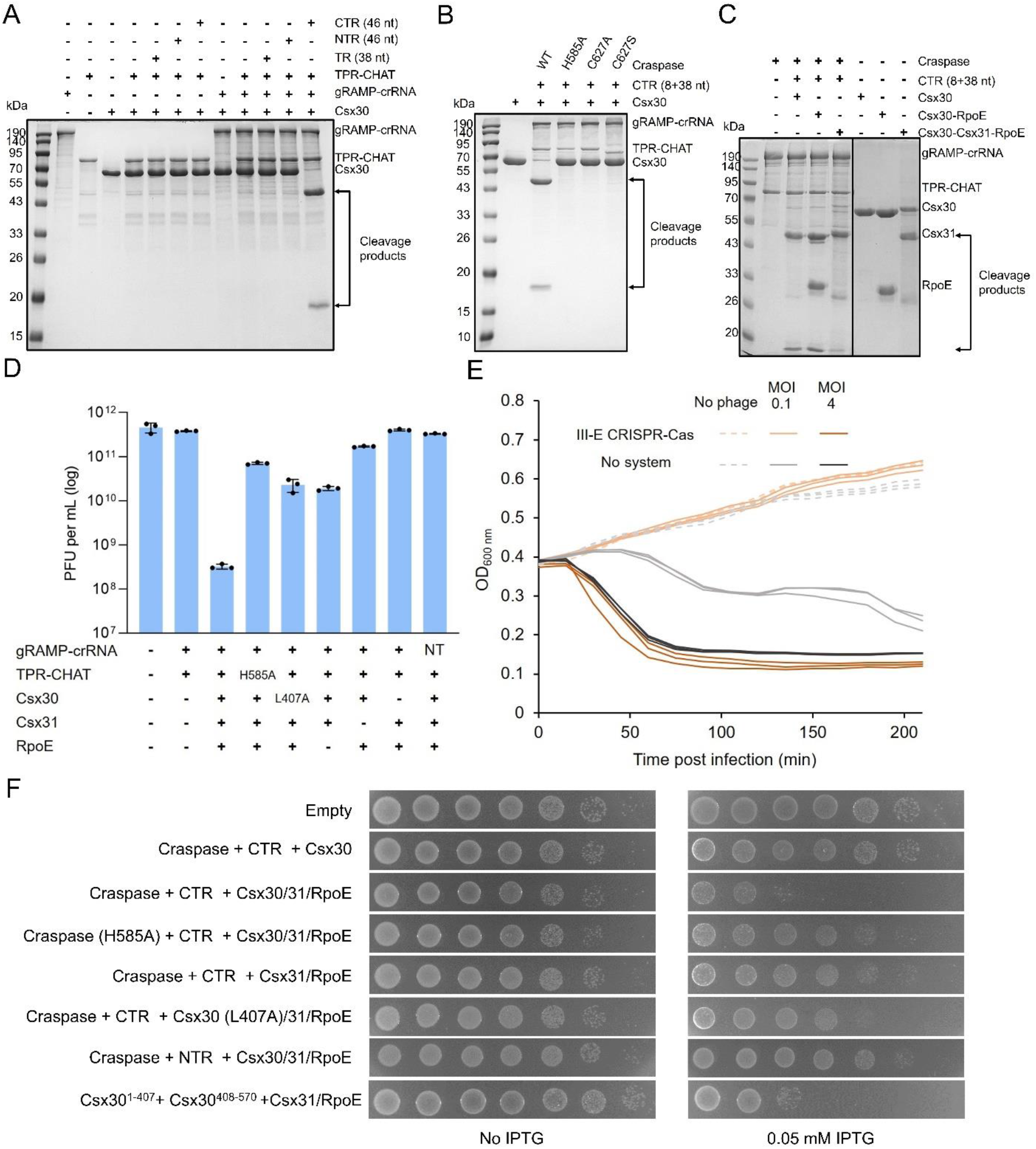
Immunity mechanism of type III-E CRISPR-Cas system. (A) The target RNA dependent Csx30 cleavage by the TPR-CHAT in the Craspase complex. The TR/NTR/CTR RNA molecules, TPR-CHAT, gRAMP-crRNA and Csx30 were added at 0.5, 0.3, 0.3 and 1.5 μM. (B-C) The target RNA dependent Csx30 cleavage with the Craspase or its mutants with mutations at the active sites of TPR-CHAT. The purified Craspase, CTR and Csx30 or Csx30-RpoE or Csx30-31-RpoE complex were added at 0.3, 0.5 and 1.5 μM. (D) The type III-E CRISPR-Cas operon protects against phages. The efficiency of plating of λ phages on *E. coli* BL21 (DE3) cells expressing genes of the type III-E CRISPR-Cas operon from *Candidatus* “Scalindua brodae” are shown. Data represent plaque-forming units (PFU) per milliliter and are the averages of three independent replicates, with individual data points overlaid. NT, non-target. (E) Growth of liquid cultures of *E. coli* cells expressing the five genes of the type III-E CRISPR-Cas operon or control *E. coli* cells (no system). Cells were infected with λ phage at 37 °C. Bacteria were infected at time 0 at an MOI of 4 or 0.1. Three independent replicates are shown for each MOI, and each curve represents an individual replicate. (F) Activation of the type III-E CRISPR-Cas system is toxic. Cells encoding the indicated plasmids were plated in 10-fold serial dilution on LB-agar in conditions without induction or induce expression (0.05 mM IPTG).

## Cleavage of Csx30 triggers an abortive infection requiring RpoE and Csx31

Next, we investigated the biological implications for the CTR-activated Csx30 cleavage. Notably, we found that Csx30 can form a stable complex with either RpoE (Fig. S8E), or RpoE-Csx31 when the three proteins are expressed together (Fig. S8F). Notably, both the Csx30-RpoE and Csx30-Csx31-RpoE complex can also act as the substrate of the Craspase (Figure 4C). After cleavage after L407 of Csx30, interestingly, the N-terminal part, but not the C-terminal part, can still bind RpoE and RpoE-Csx31 (Fig. S8G and S8H). Then, we investigated the defense mechanism of the type III-E CRISPR-Cas system. We first incorporated the genes encoding gRAMP, TPR-CHAT, Csx30, Csx31 and RpoE, and the synthetic CRISPR array containing spacers directed to the transcribed strand of early gene transcripts of the dsDNA phage lambda (λ) into *E. coli* BL21. Efficiency of Plating (EOP) assays showed that co-transforming five genes with the synthetic CRISPR array provided robust defense against phage λ. Interestingly, deletion of any of Csx30/Csx31/RpoE or using a CRISPR array with non-targeting guide abolished or decreased the protection to different extents (Figure 4D). Notably, mutation of the active site of TPR-CHAT or the cleavage site of Csx30 also significantly impaired defense, suggesting that the protease activity of TPR-CHAT towards Csx30 plays a key role for this anti-phage effect. We next investigated the anti-phage strategy of this type III-E CRISPR-Cas system through bacterial growth assays (Figure 4E). When infected in liquid media, the five-gene-containing bacterial cultures collapsed in high multiplicity of infection (MOI) like control cells but survived in low MOI infection. This indicates that the immunity mechanism of type III-E CRISPR-Cas is abortive infection, as seen in canonical type III CRISPR-Cas systems. Based on this, we speculated that cleavage of Csx30 triggered by CTR recognition by the Craspase in the presence of RpoE and Csx31 may be toxic for cells. To test this, we co-expressed the type III-E CRISPR-Cas system with a CTR sequence in *E. coli* cells, which induced potent cellular toxicity in the absence of phage infection (Figure 4F). However, H585A mutation of TPR-CHAT, L407A mutation of Csx30, deletion of Csx30 or Csx31/RpoE, or co-expressing with the NTR all decreased the cellular toxicity to different extents (Figure 4F). Notably, expression of the two segments of Csx30 with Csx31 and RpoE directly cause cell toxicity (Figure 4F, bottom lane). Taken together, our results showed that CTR-induced cleavage of Csx30 by the Craspase triggers abortive infection which requires RpoE and Csx31.

## Potential mechanism of the toxicity induced by Csx30 cleavage

Next, we investigated the mechanism of cell toxicity induced by Csx30 cleavage in the presence of Csx31 and RpoE. First, the structure of Csx30, Csx31, RpoE and Csx30-RpoE were predicted using AlphaFold2 (Fig. S9 A-D) (30). The Csx30 comprises an N-terminal domain (NTD) and C-terminal domain (CTD. Notably, the cleavage site L407 is located at a β-hairpin in the CTD (Fig. S9A). The structure of Csx31 also displays two domains, NTD and CTD (Fig. S9B). Dali search with Csx30 and Csx31 do not return entries with a high sequence coverage, giving limited functional insights. The structure of RpoE displays a typical fold of sigma factor showing homology to multiple transcriptional regulators (Fig. S9C). Consistent with the purified Csx30-RpoE complex, the predicted Csx30-RpoE complex displays an extensive binding interface (Fig. S9D) with a buried area of 3773.8 Å^2^. The location of the L407 of Csx30 also supports that Csx30^1-407^ still interacts with RpoE and Csx30^408-570^ does not (Fig. S9D). Interestingly, prediction of the Csx30-Csx31-RpoE structure does not return credible results (with severe steric clash) with the full-length Csx30 (Fig. S9E). However, when Csx30^1-407^ was used, the three proteins can form a complex (Fig. S9F), in which Csx31 partially overlaps with the position of Csx30^408-570^ in the Csx30-RpoE complex, suggesting the structural flexibility of the linker between Csx30 NTD and CTD. Based on these structure prediction results, we propose that the Csx30^1-407^ may induce cell toxicity as a complex with Csx31 and RpoE, or this ternary complex may undergo other changes to disassembly to elicit cell death.

## Structures of Craspase bound with Non-cognate Target RNA molecules

To investigate how the protease activity of the Craspase is triggered by CTR but not TR or NTR binding, we first determined the cryo-EM structure of Craspase bound to TR at 2.97 Å resolution (Figure 5A; Fig. S1; and Table S1). The TR adopts a similar structure as in the gRAMP-crRNA-TR structure and does not directly interact with the TPR-CHAT protein (Figure 5A). Structural alignment between the Craspase and Craspase-TR also reveals an overall similarity, with an RMSD of 0.50 Å among 1550 Cα atoms. Like in the context of the gRAMP-crRNA, only the Cas11 domain of gRAMP displays a marked rotation (Figure 5B and 5C). TR binding also causes slight conformational change in the N-terminal two helices in the TPR domain (Fig. S10A). However, no apparent rotation or translation was observed for the entire TPR-CHAT upon TR binding, compared to the apo Craspase structure (Figure 5B). To further determine how the 3’ anti-tag region of non-cognate target RNA interacts with the Craspase, we solved the cryo-EM structure of Craspase complexed with NTR, which is longer than TR by a further 8-nt sequence complementary to the 5’ tag region of crRNA (Figure 5D and 5E; Fig. S1; and Table S1). The gRAMP D547A/D698A mutant was used to obtain the complex to prevent target RNA cleavage. Then we aligned the structure of the Craspase and Craspase-NTR at the gRAMP part (Fig. 5F). Notably, a rearrangement of TPR-CHAT is observed between the apo and NTR structures. The TPR-CHAT rotates by ~3.6° and translates ~0.3 Å around the axis, and displays a ~0.7 Å movement of its mass center. This is primarily a rigid body movement of TPR-CHAT, because TPR-CHAT alone in the two structures align well except slight rotations of its N-terminal two helices (Fig. S10B). Comparison between Craspase bound with TR and NTR showed that binding of the 3’ anti-tag of NTR further rearranges the Cas7.2 loop (G376-D386) and TPR-CHAT, and meanwhile a loop within the Cas7.2 domain (residues K443-N453) becomes disordered upon NTR binding (Figure 5G). While the NTR used in this study contains eight nucleotides in the 3’ anti-tag, the density only allows us to model six of them ((−1)’-(−6)’, Figure 5H). Among the six nucleotides in the NTR, only nucleotides (−1)’ and (−2)’ form two base pairs with nucleotides (−1) and (−2) within the 5’ tag of crRNA (Figure 5I). A(−3)’ is also kinked by the “thumb” of Cas7.1 domain, flips out like U(−3) in the crRNA, and is sandwiched by the loop (L384-Y389) of Cas7.2 and the TPR domain of TPR-CHAT. For the upstream nucleotides C(−4)’-G(−6)’, interestingly, a loop in the Cas7.2 domain (G488-C503) protrudes into the crRNA-NTR duplex at this region, thus preventing base paring at these nucleotides (Figure 5J).

**Figure 5.**
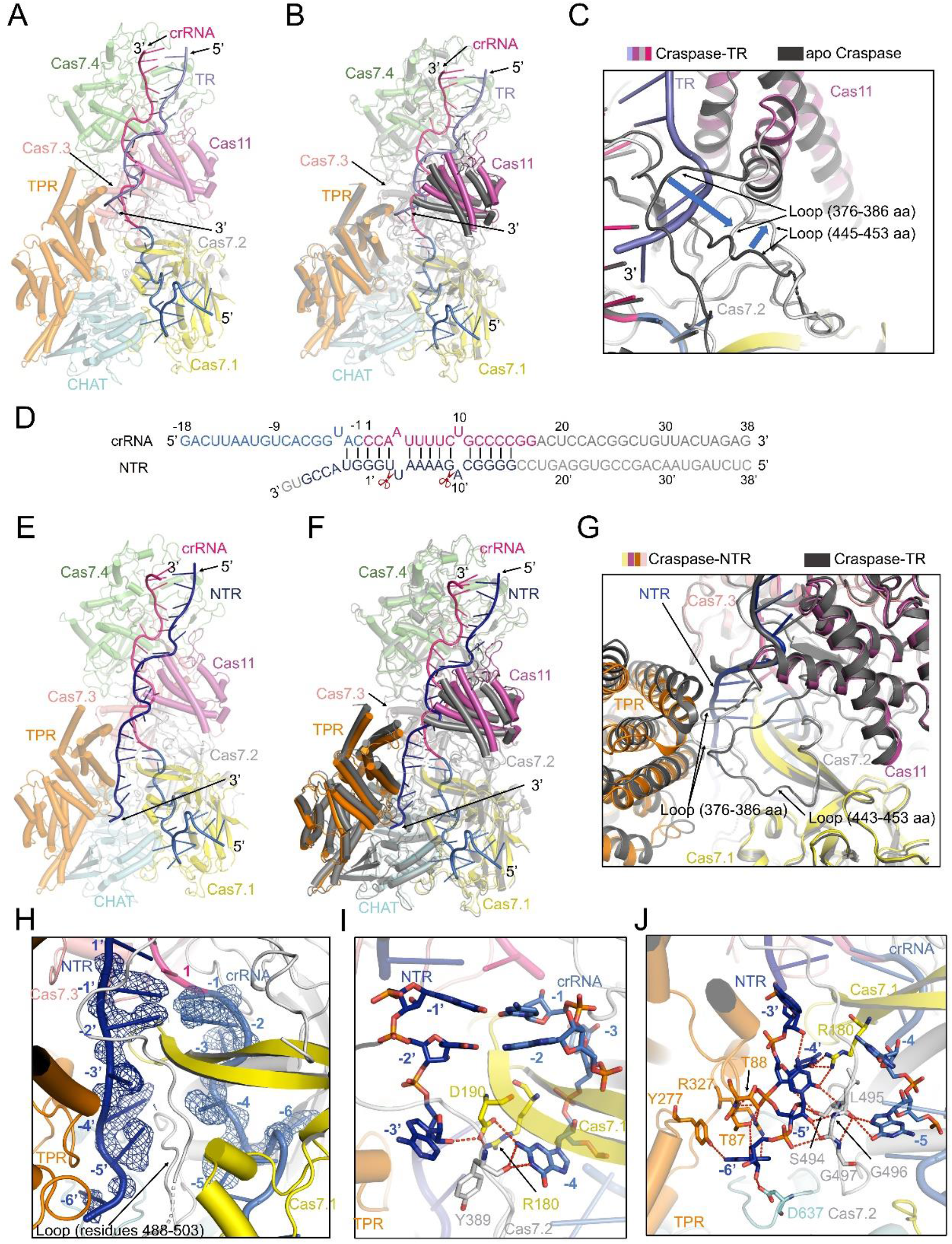
Cryo-EM structures of the Craspase-TR and Craspase-NTR. (A) Overall structure of the Craspase (colored as in Figure 3A) bound with TR (colored slate). (B) Structural superimposition of the Craspase (colored dark gray) and Craspase-TR (colored as in A). (C) Close-up view of the two loops adjacent to the Cas11 domain in the structural superimposition in B. The Craspase and Craspase-TR are colored as in B. (D) Schematic representation of the crRNA-NTR duplex in the Craspase-NTR. The nucleotides of crRNA and NTR used in the experiment are depicted in lines. The NTR is colored in dark blue. The invisible nucleotides are colored in gray. (E) Overall structure of the Craspase (colored as in Figure 3A) bound with NTR (dark blue). (F) Structural superimposition of the Craspase (colored dark gray) and Craspase-NTR (colored as in E). (G) Close-up view of the region of the two indicated loops in the structural superimposition between the Craspase-NTR (colored as in E) and Craspase-TR (colored dark gray). Note the disordered loop (residues 443-453) in the Craspase-NTR structure. (H) Six nucleotides could be modelled in the 3’ anti-tag region of NTR. The Craspase-NTR is colored as in E. The densities corresponding to (−1)’-(−6)’ of NTR and (−1)-(−6) of crRNA are shown in mesh. (I-J) Detailed interactions between the Craspase and (−1)’-(−3)’ of NTR (I) and (−4)’-(−6)’ of NTR (J).

## Overall Structure of the Cognate Target RNA-Bound Craspase Complex

Then, we moved on to determine the cryo-EM structure of the Craspase (gRAMP D547A/D698A) bound to CTR at 3.27 Å resolution (Figure 6A and 6B; Fig. S1; and Table S1). In the Craspase-CTR structure, the spacer region forms base pairs similar to what is seen in the TR- and NTR-bound Craspase complex. However, in the structure, the 3’ anti-tag of CTR displays a strikingly different conformation and binding channel in the TPR domain, compared with those of NTR in the Craspase-NTR structure (Figure 6C, or compare Figure 6B and 5E). In the Craspase-CTR structure, TPR-CHAT is also rotated and translated around the gRAMP axis with respect to the apo structure, as observed in the NTR bound complex. However, the TPR-CHAT of the CTR-bound structure is rotated ~5.2° and translates ~1.2 Å, and displays a ~2.8 Å movement of its mass center, suggesting that lack of base pairing between the 5’ tag and 3’ anti-tag, together with the extensive interactions between the 3’ anti-tag of CTR and TPR-CHAT promoted a further step of ~1.6° and ~0.9 Å (and ~1.1 Å in its mass center) of TPR-CHAT in the CTR-bound with respect to the NTR-bound Craspase (Figure 6C).

**Figure 6.**
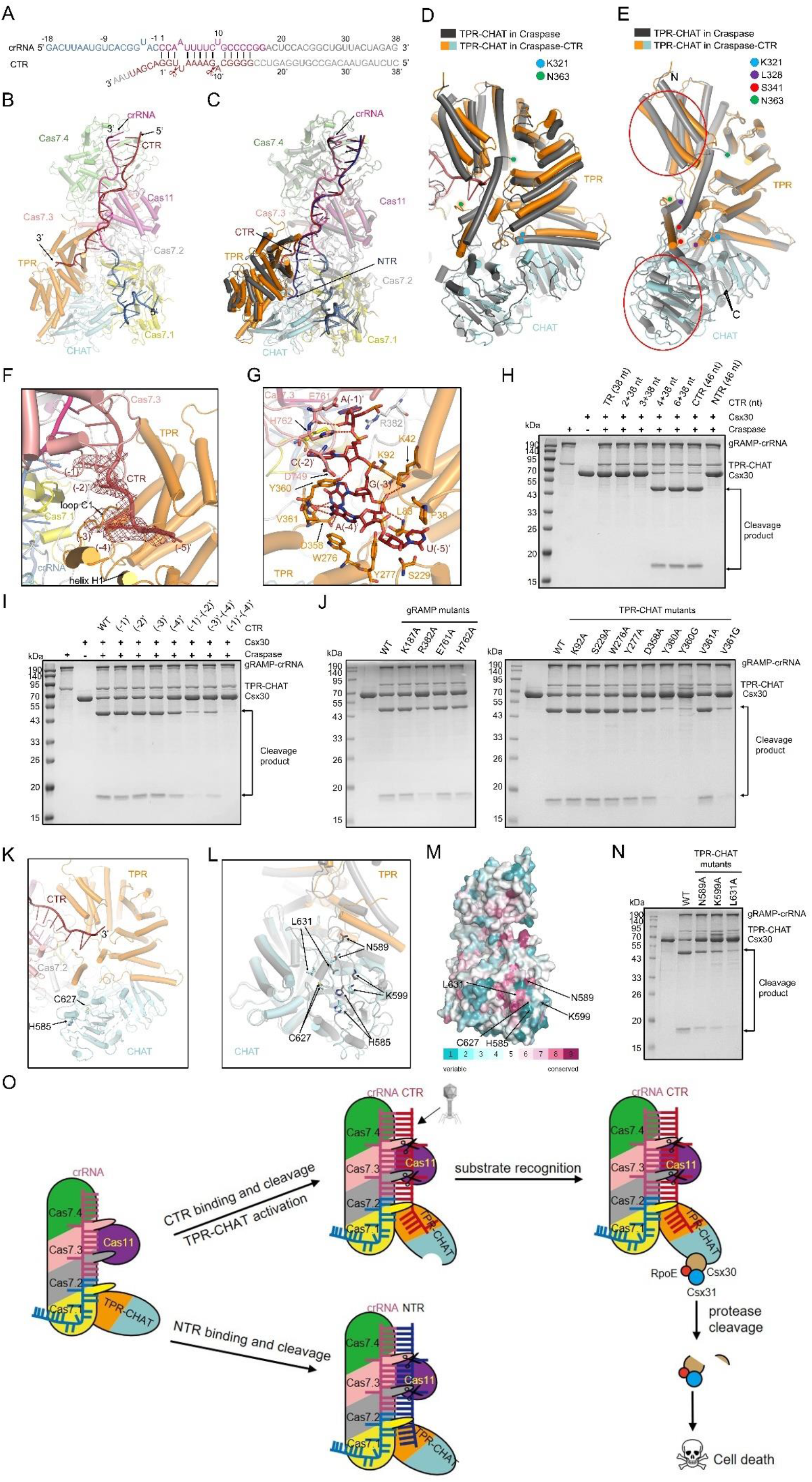
Cryo-EM structure of the Craspase-CTR and CTR-recognition mechanism. (A) Schematic representation of the crRNA-CTR duplex in the Craspase-CTR. The nucleotides of crRNA and CTR used in the experiment are depicted in lines. The CTR is colored in firebrick. The invisible nucleotides are colored in gray. (B) Overall structure of the Craspase (colored as in Figure 3A) bound with CTR (firebrick). (C) Structural superimposition of the Craspase-CTR (colored as in B) and Craspase-NTR. In the Craspase-NTR structure, NTR is colored dark blue and the other regions are colored dark gray. (D) Structural alignment of apo Craspase (colored dark gray) and Craspase-CTR (colored as in B) at the gRAMP part and the close-up view of the TPR-CHAT region is shown. The positions corresponding to K321 and N363 in the two structures are marked with colored circles. (E) Structural alignment of TPR-CHAT in the apo Craspase (colored dark gray) and Craspase-CTR (colored as in B). The positions corresponding to K321, L328, S341 and N363 in the two structures are marked with colored circles. Apart from the helix (D339-V361), subregions in the two domains of TPR-CHAT with notable conformational changes are marked with red circles. (F) Five nucleotides could be modelled in the 3’ anti-tag region of CTR. The Craspase-CTR is colored as in B. The densities corresponding to (−1)’-(−5)’ of CTR are shown in mesh. (G) Detailed interactions between the Craspase and (−1)’-(−5)’ of CTR. (H) The target RNA dependent Csx30 cleavage with the Craspase and CTR molecules with different numbers of mismatch nucleotides in the 3’ anti-tag region. The purified Craspase, Csx30 and different RNA molecules were added at 0.1, 1.5 and 0.25 μM, respectively. (I) The target RNA dependent Csx30 cleavage assays showing the impact of 3’ anti-tag mutation on the protease activity of the Craspase. The nucleotide numbers in the fist line represent the sites mutated to crRNA complementary nucleotides. The purified Craspase, Csx30 and CTR mutants were added at 0.1, 1.5 and 0.25 μM, respectively. (J) The target RNA dependent Csx30 cleavage assays showing the impact of gRAMP (left panel) and TPR-CHAT mutations (right panel) to hinder the recognition of the 3’ anti-tag of CTR. The gRAMP-crRNA, TPR-CHAT, Csx30 and CTR were added at 0.1, 0.1, 1.5 and 0.25 μM, respectively. (K) Cartoon representation of the binding site of the 3’ anti-tag region of CTR and protease active site in TPR-CHAT. H585 and C627 of CHAT domain are shown in sphere model. (L) Allosterically rearrangement of the active site of CHAT domain after CTR binding. TPR-CHAT in the CTR-bound Craspase is colored in orange and cyan. TPR-CHAT in the apo Craspase in colored in gray. Active site residues and several surrounding residues are shown as sticks. (M) Sequence conservation through ConSurf analysis mapped to the surface of TPR-CHAT. The color scheme describes the degree of conservation as indicated. H585, C627 and several conserved surrounding residues of TPR-CHAT are shown as sticks. (N) The target RNA dependent Csx30 cleavage assays showing the impact of TPR-CHAT mutations to possibly hinder Csx30 recognition. The gRAMP-crRNA, TPR-CHAT, Csx30 and CTR were added at 0.1, 0.1, 1.5 and 0.25 μM, respectively. (O) Model of type III-E CRISPR-Cas immunity. The Craspase complex recognizes target RNA which contains sequence complementary to the crRNA, by forming the crRNA-target RNA duplex. The target RNA is cleaved at 6-nt intervals by the Cas7.2 and 7.3 domains, respectively. The 3’ anti-tag region of CTR binds to and induces conformational changes of TPR-CHAT, which allosterically activates the protease activity of TPR-CHAT. Activated TPR-CHAT then cleaves Csx30, which together with RpoE and Csx31 induces abortive infection.

## CTR binding causes dramatic conformational change in TPR-CHAT

Apart from rotation and translation of TPR-CHAT, more importantly, binding of CTR results in marked conformational changes in TPR-CHAT (Figure 6D), which is clearly different from its simply rigid-body/no movement caused by NTR/TR binding. This is in agreement with the result that only CTR can activates the protease activity of TPR-CHAT (Figure 4A). The most dramatic conformational change happens in the K321-N363 region of the TPR domain (Figure 6D). In the apo (or TR/NTR-bound) Craspase, this region forms a loop-helix (L328-K333)-loop-helix (D339-V361) structure. However, upon CTR binding, this region turns to a helix (F324-R327)-X-helix (N342-Q356)-loop (Q357-363, named loop C1) structure, in which “X” represents the lack of density for S329-E340, indicating flexibility of this region in CTR-bound Craspase (Figure 6E). Notably, the 23-residue helix (D339-V361) in the apo Craspase displays a ~43° rotation towards the CHAT domain and shortens to a 15-residue (N342-Y356) helix upon CTR binding (Figure 6E). The extreme C-terminal residues of TPR domain fold as a short loop (loop C1) to interact with CTR (Figure 6F and 6G). Meanwhile, the loop (E268-R280) between the helix (W251-E268) and the helix (H281-D297) also swings away from the CHAT domain and forms a helix (T271-W278, named helix H1) to bind the 3’ anti-tag of CTR upon its binding (Figure 6F and 6G). In addition to these marked rearrangements, parts of both TPR (residues 14-115) and CHAT (residues 527-653) domain also display notable conformational changes (Figure 6E). Taken together, CTR mainly binds the loop C1 and helix H1 in the TPR domain of TPR-CHAT, and triggers dramatic conformational changes in both TPR and CHAT domains.

## Detailed interactions between 3’ Anti-tag of CTR and Craspase

While the CTR used in this study contains eight nucleotides in the 3’ anti-tag, the density only allows us to model five of them ((−1)’-(−5)’) (Figure 6F). The phosphate-ribose backbone of the 3’ anti-tag is stabilized by hydrogen bonds from K42, L83 and Y360 of TPR-CHAT and E761 and H762 of gRAMP (Figure 6G). Moreover, sidechains of Y360 and W276 also form stacking interactions with C(−2)’ and A(−4)’, respectively. Notably, U(−5) does not form hydrogen bonds with the Craspase and only a hydrophobic interaction with P38 of TPR-CHAT. Then, we investigated the minimal numbers of mismatch nucleotides in the CTR required to activate the protease activity of Craspase. To this end, we synthesized different CTR molecules with 2-8 nt mismatches in their 3’ ends and tested their ability of Craspase activation in the protease assay. The results showed that CTR with as short as 4 nt mismatches can still activate the protease activity of Craspase, but those with fewer than 4 nt mismatches cannot, suggesting that 4 nt mismatches in the 3’ end are the minimal requirement for CTR (Figure 6H). Moreover, to evaluate the impact of the degree of non-complementarity of the 3’ anti-tag, we measured the effects of mutations of the (−1)’-(−4)’ nucleotides within the 3’ anti-tag of the CTR (with 8 nt mismatches) on the protease activity of Craspase (Figure 6I). For single nucleotide mutations, only A(−4)’ in CTR reduced Csx30 cleavage in the protease activity assay. The two double mutations at (−1)’-(−2)’ and (−3)’-(−4)’ resulted in greater reduction of protease activity (Figure 6I). Mutation of (1)’-(−4)’ simultaneously abolished the protease activity. In turn, mutations of the Craspase residues involved in 3’ anti-tag binding showed that R382A and H762A of gRAMP, and D358A, Y360A/G and V361G of TPR-CHAT reduced the protease activity activated by CTR (Figure 6J). Together, these results confirm that non-complementarity between nucleotides (−1)’-(−4)’ of the 3’ anti-tag within target RNA and the 5’ tag of the crRNA is critical for activating the protease activity of the Craspase.

## Mechanism of activation of CHAT activity

The binding channel of CTR in the TPR domain is away from the catalytic sites H585 and C627 of the CHAT domain (Figure 6K), but CTR binding also causes marked conformational changes in the CHAT domain (Figure 6D). That is, CTR allosterically activates the peptidase activity of TPR-CHAT. Notably, structural alignment of TPR-CHAT within the apo and CTR-bound Craspase showed marked differences in the relative positions of the catalytic sites H585 and C627 and their surrounding residues (Figure 6L). As the catalytic His and Cys residues are directly involved in the hydrolysis of the scissile peptide bond, their relative positions in the enzymatic active site are important for the peptidase activity (31). Moreover, the residues surrounding H585 and C627 on the same surface are also largely conserved (Figure 6M). We hypothesized that these residues may be involved in substrate recognition. Craspase mutants containing mutations of several conserved residues on this surface displayed reduced CTR-activated protease activities (Figure 6N), supporting our hypothesis. Taken together, we propose a model in which TPR-CHAT in the Craspase is itself in an inactivated state, and CTR binding-induced conformational change in TPR-CHAT in the context of the Craspase rearranges the active site to be suitable for recognition and cleavage of the Csx30 substrate, which induces abortive infection in the presence of Csx31 and RpoE (Figure 6O).

## Discussion

Different from other canonical type III CRISPR-Cas systems, the type III-E CRISPR-Cas system does not contain the signature Cas10 subunit, thus lacks the activities of ssDNA cleavage and cOA synthesis. Our study uncovers the structural and functional characteristics of gRAMP and the Craspase, and reveals the mechanism of type III-E CRISPR-Cas immunity. Our data show that Craspase is the intact form of the type III-E CRISPR-Cas effector which contains both target RNA cleavage and protease activity. Importantly, the protease activity of Craspase towards Csx30 is controlled by complementarity between the 5’ tag of crRNA and the 3’ anti-tag of target RNA. Our structural analysis of Craspase complexes bound to different targets reveals that nucleotides (−1)’-(−4)’ of 3’ anti-tag are essential for activating the protease activities, thus providing the structural basis for control of target RNA-dependent Csx30 cleavage. Moreover, we also prove that CTR-binding-induced cleavage of Csx30 induces an abortive infection in the presence of Csx31 and RpoE, as the antiviral strategy of the type III-E CRISPR-Cas system.

In this study, we identified two aspartate residues D698/D806 of *Sb*-gRAMP responsible for target RNA cleavage at Site 2 and one aspartate residue D547 responsible for cleavage at Site 1. Mutation of S457, the corresponding residue of D698 in the canonical “catalytic loop” of Cas7.2 does not interfere with RNA cleavage. Interestingly, in the corresponding position of *Sb*-gRAMP S457, serine is replaced by aspartate or glutamate residues in some gRAMP homologs (Fig. S3). For example, in the *Di*-gRAMP, the corresponding residue D429 has been shown to be responsible for Site 1 cleavage (17). Meanwhile, D547 of *Sb*-gRAMP is replaced by N518 in the corresponding position in *Di*-gRAMP. However, in gRAMP from *Deferribacteres* bacterium, at the two positions are E401 and D492, respectively, suggesting possible cleavage activity (Fig. S3). Taken together, these data suggest that each cleavage site of target RNA might be cleaved by two aspartate/glutamate residues in the catalytic loop and “thumb” subdomain, respectively, or only one aspartate/glutamate residue at either of the two regions. Future studies could test this hypothesis in other gRAMP orthologs as well as other type III CRISPR-Cas systems.

The mechanisms of type III-A/B CRISPR-Cas immunities have been investigated by extensive studies. However, no immunity mechanism has been revealed for type III-E CRISPR-Cas system before our study. van Beljouw et al. proposed that target RNA recognition might activate the protease activity of the Craspase to confer viral immunity, however, no such activity or viral immunity mechanism has been uncovered by their study and other studies (16). In addition, Özcan et al. showed that TPR-CHAT displays weak inhibition on target RNA cleavage by the *Di*-gRAMP *in vitro* (17), but the *in vitro* study by van Beljouw et al. did not show such inhibition by TPR-CHAT using *Sb*-gRAMP (16). Our structural and functional studies uncovered that the Craspase is the intact Cas effector conferring viral immunity of type III-E system, which discriminates foreign nucleic acids through non-complementarity between the 5’ tag of crRNA and 3’ anti-tag of RNA target. CTR binding to the Craspase activates the protease activity of TPR-CHAT to cleave Csx30, which induces an abortive infection in the presence of Csx31 and RpoE. Interestingly, van Beljouw et al. and Özcan et al. both performed *in vivo* assays of the type III-E system in their studies but did not uncover the immunity mechanism of this system (16,17). In the study by van Beljouw et al., they co-expressed the Craspase with an inducible target RNA in *E. coli* and did not observe growth defects upon target RNA production. Lack of co-expression of Csx30, Csx31 and RpoE may explain their results. In the study by Özcan et al., they found that a single *Di*-gRAMP gene with targeting crRNA is sufficient for MS2 RNA phage interference in an *E. coli* system, and TPR-CHAT even inhibits the interference activity of *Di*-gRAMP *in vivo* (17). Our explanation for the discrepancy between their and our studies is as follows. DNA phage λ and RNA phage MS2 were used in our and their phage interference studies, respectively. Thus, one possible reason is that for RNA phages, the invading RNA could be targeted by the gRAMP-crRNA upon their invasion. Therefore, a single gRAMP gene with targeting crRNA can provide sufficient phage interference by cleaving the invading RNA. However, for DNA phages, the type III-E immunity requires transcription of phage DNA, which produces target nascent RNA molecules. In this case, simply cleaving transcribed RNAs might be insufficient to interfere with phage propagation, therefore, abortive infection through activation of the protease activity of TPR-CHAT is adopted by the infected cell. Of course, another possibility is that the type III-E systems from different species may adopt distinct immunity mechanisms. Interestingly, several type III-E system-containing species lack one or two of Csx30/Csx31/RpoE or even all of them (16,17), future studies should be conducted to investigate the similarities and differences of their immunity mechanisms.

It still remains not fully understood how cleavage of Csx30 triggers abortive infection in the presence of Csx31 and RpoE. Interestingly, divided at the cleavage site, while the N-terminal part of Csx30 is largely conserved among its homologs, the C-terminal part and even the cleavage site L407 is not conserved (Fig. S11). This suggests that after cleavage by the Craspase, the N-terminal part of Csx30 might play a more important role than the C-terminal part. For the cleavage site, it has been known that proteases of the caspase family typically display aspartate P1 cleavage specificity, while those of metacaspases and paracaspases cleave substrates after an arginine or lysine residue (32). Coincidentally, a very recent study discovered that bacterial gasdermin (bGSDM) homologs also defended against phages and executed cell death. They were activated by site-specific cleavage catalyzed by dedicated caspase-like proteases, which removed an inhibitory C-terminal peptide from bGSDMs (33). It was found that the *Runella* bGSDM cleavage site also occurs after a P1 leucine residue, L247. This cleavage residue is not conserved either, as a *Bacteroidetes* bGSDM (metagenomic scaffold) was cleaved after a P1 residue arginine 247 (33). More interestingly, another recent study found that, an effector protein predicted to be linked to type III CRISPR-Cas systems contains a SAVED and a Lon protease domain, which upon activation by cA4 cleaves a CRISPR-T protein (34). The releasing fragment of CRISPR-T is structurally similar to MazF toxins and predicted to be a sequence specific nuclease. Neither of Csx30 and Csx31 has any identifiable domains or predicted molecular functions using BLASTp and HHPred, or AlphaFold2 and DALI servers, and only RpoE displays sequence similarity to the sigma24 transcription factor. Notably, both the full-length and the N-terminal part of Csx30 (residues 1-407) stably bind RpoE and RpoE-Csx31. Very recently, a preprint by Nishimasu and coworkers suggested that the N-terminal of Csx30 is toxic to cells using the type III-E system from *Desulfonema ishimotonii (35)*, however the detailed mechanism still awaits investigating especially in the context of both Csx31 and RpoE. Taken together, our study reveals both the catalytic mechanism of the gRAMP and the unprecedented immunity mechanism of the type III-E CRISPR-Cas system.

## Acknowledgements

We thank Prof. Jianlin Lei and Fan Yang (Tsinghua University), Prof. Peiyi Wang and Xiaomin Ma (Southern University of Science and Technology) for EM data collection. We thank the Tsinghua University Branch of the China National Center for Protein Sciences (Beijing) and Southern University of Science and Technology for providing the cryo-EM facility support. We would like to thank Mrs. Wu Yao (State Key Laboratory of Plant Genomics, Institute of Microbiology, Chinese Academy of Sciences) for assistance in the microscale thermophoresis (MST) experiment. This work was supported by the National key research and development program of China (2017YFA0506500, 2019YFC1200500 and 2019YFC1200502), National Natural Science Foundation of China (32171274 and 32000901), Beijing Nova program and the Fundamental Research Funds for the Central Universities (XK1802-8).

## Author Contributions

Y. F. conceived and supervised the project. X. L., H. W., Y. X., L. H., Z. G., N. L., F. L. and W. X. purified the proteins, prepared the protein-nucleic acid complexes, performed the activity analysis and binding assays supervised by Y. F. and Y. Z.. X. L. and Z. G. performed phage assays. L. Z. prepared the cryo-EM sample, collected the cryo-EM data and solved the cryo-EM structures supervised by M. Y.. T. G. conducted preliminary structure prediction. Y. F. analyzed the data and wrote the paper with assistance from all the authors.

## Materials and Methods

### Protein expression and purification

The full-length gRAMP, TPR-CHAT, Csx31 and RpoE genes were synthesized by GenScript, amplified by PCR, and cloned into pET28a to produce His_6_-SUMO-tagged fusion proteins with a SUMO-specific protease cleavage site between His_6_-SUMO and the target protein, respectively, and the crRNA fragment were cloned into pACYCDuet-1. The full-length Csx30 genes were synthesized by GenScript and amplified by PCR and cloned into pGEX6p-1 to produce GST-tagged fusion proteins with a PreScission Protease cleavage site between GST and the target protein. The gRAMP-crRNA complex was generated through co-expression of the two plasmids in *E. coli* strain BL21. The Craspase complex was generated through co-lysed the gRAMP-crRNA complex with TPR-CHAT cloned into pGEX6p-1. For the Csx30-RpoE complex, the full-length Csx30 or Csx30^1-407^ were cloned into pRSFDuet-2 respectively, the RpoE was cloned into pRSFDuet-1 to produce His_6_ -tagged fusion proteins and expressed in *E. coli* strain BL21.

The proteins were expressed in *E. coli* strain BL21 and induced by 0.2 mM isopropyl-β-D-thiogalactopyranoside (IPTG) when the cell density reached an OD_600nm_ of 0.8. For His_6_-SUMO-tagged protein, the cells of the gRAMP-crRNA complex and the Craspase complex were harvested, re-suspended in lysis buffer (50 mM Tris-HCl pH 8.0, 500 mM NaCl, 30 mM imidazole) and lysed by sonication. The cell lysate was centrifuged at 13,000 rpm for 50 min at 4 °C to remove cell debris. The supernatant was purified through Ni-column, the fusion protein was then digested 2.5 h with the SUMO-specific protease at 18 °C. The eluant was concentrated and further purified by Heparin chromatography (the TPR-CHAT was purified by anion exchange chromatography) and Superdex-200 (GE Healthcare) column equilibrated with a buffer containing 10 mM Tris–HCl pH 8.0, 500 mM NaCl, 5 mM DTT and 5% glycerol. The purified protein was analyzed by SDS-PAGE.

For GST-tagged protein, after induction, the cells were harvested, re-suspended in lysis buffer (1×PBS, 2 mM DTT and 1 mM PMSF) and lysed by sonication. The cell lysate was centrifuged at 13,000 rpm for 50 min at 4 °C to remove cell debris. The supernatant was applied onto a self-packaged GST-affinity column (2 mL Glutathione Sepharose 4B; GE Healthcare) and contaminant proteins were removed with wash buffer (20 mM HEPES pH 7.5, 200 mM NaCl, 2 mM DTT). The fusion protein was then digested 2h with PreScission protease at 18 °C. The eluant was concentrated and further purified using a Superdex-200 (GE Healthcare) column equilibrated with a buffer containing 10 mM Tris–HCl pH 8.0, 200 mM NaCl, and 5 mM DTT.

All mutants were generated by two-step PCR and were subcloned, overexpressed and purified in the same way as wild-type protein.

### *In vitro* RNA cleavage assay

For testing the activity of gRAMP-crRNA and their mutants, RNA cleavage reactions were performed in a 20 μL reaction volume containing purified 300 nM Sb-gRAMP-crRNA complex, 500 nM labelled RNA oligo with 5’-FAM, 11 mM Tris-HCl, 67 mM NaCl, 4.5 mM DTT and 1 mM MgCl_2_ unless stated otherwise. Typical reactions were incubated at 20 °C for 15 minutes and quenched with 5 μL the mixture of 5% SDS and 0.25 M EDTA and 100 °C for 5 minutes to stop the reactions. The products were separated by electrophoresis over 14% polyacrylamide gels containing 7 M urea and visualized by fluorescence imaging. (pre-run at 600 V for 1 hour, sample run at 250 V for 30 minutes).

Target RNA sequence (36 nt: 8 nt non-matching PFS+28 nt spacer) AGCCGUGGAGUCCGGGGCAGAAAAUUGGACGAUUAA

### Mass spectrometry

LC-MS was used to determine the intact molecular weight of purified full-length Csx30 and its two fragment products obtained from the protease cleavage. For LC-MS analysis, the analytes were separated by a 10 min gradient elution at a flow rate 0.5 mL/min with an ACQUITY UPLC system, which was directly interfaced with a SYNAPT-G2-Si mass spectrometer produced by Waters company. The analytical column was a Protein BEH C4 silica capillary column (2.1mm ID, 50 mm length; Made in Ireland) packed with C-4 resin (300 Å, 1.7 μm) purchased from Waters company. Mobile phase A consisted of 0.1% formic acid aqueous solution, and mobile phase B consisted of 100% acetonitrile and 0.1% formic acid.

Aliquots of 3 μL analytes were loaded into an autosampler for electrospray ionization. Samples were analyzed on a Q-TOF mass spectrometer (SYNAPT G2-Si, Waters company) instrument optimized for high-mass protein analysis. The measurements were performed with capillary 3000–3500 V and data were collected over the *m/z* range of 500–2000. Once having acquired raw electrospray mass spectra, the raw spectrum can be deconvoluted by MaxEnt 1 (Waters) to generate a spectrum (relative intensity versus mass) where all the charge-state peaks of a single species have been collapsed into a single (zero-charge) peak.

### Protease cleavage assays

The TPR-CHAT cleavage reactions were typically assembled in 20 μL volumes with buffer containing 200 mM Tris-HCl (pH 8.0), 100 mM NaCl. 0.3 μM Craspase was added to reactions with 0.5 μM CTR or NTR or TR, 1.5 μM Csx30 and incubated at 20°C for 1 h. For the mutants of TPR-CHAT, 0.1 μM gRAMP-crRNA complex and 0.1 μM TPR-CHAT protease or its mutants were added to reactions with 0.25 μM CTR, 1.5 μM Csx30 and incubated at 20°C for 30 min. Reactions were resolved by 15% SDS-PAGE, stained with Coomassie blue R-250.

CTR sequence (46 nt: 8 nt non-matching PFS+38 nt spacer) CUCUAGUAACAGCCGUGGAGUCCGGGGCAGAAAAUUGGACGAUUAA

NTR sequence (46 nt: 8 nt matching PFS+38 nt spacer) CUCUAGUAACAGCCGUGGAGUCCGGGGCAGAAAAUUGGGUACCGUG

TR sequence (38 nt: 38 nt spacer) CUCUAGUAACAGCCGUGGAGUCCGGGGCAGAAAAUUGG

2+38

CUCUAGUAACAGCCGUGGAGUCCGGGGCAGAAAAUUGGAC

3+38

CUCUAGUAACAGCCGUGGAGUCCGGGGCAGAAAAUUGGACG

4+38

CUCUAGUAACAGCCGUGGAGUCCGGGGCAGAAAAUUGGACGA

6+38

CUCUAGUAACAGCCGUGGAGUCCGGGGCAGAAAAUUGGACGAUU

(−1)’

CUCUAGUAACAGCCGUGGAGUCCGGGGCAGAAAAUUGGGCGAUUAA

(−2)’

CUCUAGUAACAGCCGUGGAGUCCGGGGCAGAAAAUUGGAUGAUUAA

(−3)’

CUCUAGUAACAGCCGUGGAGUCCGGGGCAGAAAAUUGGACAAUUAA

(−4)’

CUCUAGUAACAGCCGUGGAGUCCGGGGCAGAAAAUUGGACGCUUAA

(−1)’-(−2)’

CUCUAGUAACAGCCGUGGAGUCCGGGGCAGAAAAUUGGGUGAUUAA

(−3)’-(−4)’

CUCUAGUAACAGCCGUGGAGUCCGGGGCAGAAAAUUGGACACUUAA

(−1)’-(−4)’

CUCUAGUAACAGCCGUGGAGUCCGGGGCAGAAAAUUGGGUACUUAA

### Gel filtration assay

Protein samples along with the gel filtration standard purchased from Bio-Rad were respectively applied to a size-exclusion chromatography column (Superdex-200 increase 10/300 GL, GE Healthcare) equilibrated with a buffer containing 10 mM Tris-HCl pH 8.0, 200 mM NaCl, and 5 mM DTT. The assays were performed with a flow rate of 0.5 mL/min and an injection volume of 0.5 mL for each run. Samples taken from relevant fractions were applied to SDS-PAGE and visualized by Coomassie blue staining. TPR-CHAT was incubated with the Csx30-RpoE complex at 4 °C for 2 h with a molar ratio of 1:2. The incubation samples were analyzed as described above.

### Pre-crRNA processing

The pre-crRNA was transcribed *in vitro* using HiScribe™ T7 Quick High Yield RNA Synthesis Kit (NEW ENGLAND BioLabs). Transcription template (dsDNA) for pre-crRNA was generated by PCR with the template plasmid which harbors the CRISPR array used in protein expression. The pre-crRNA was purified using Monarch RNA Cleanup Kit (NEW ENGLAND BioLabs). RNAs were resuspended in diethyl pyrocarbonate H_2_O and stored at −80°C.

The RNA process reactions were performed in a 20 μL reaction volume containing purified 1 μM Sb-gRAMP or gRAMPΔinsertion, 500 nM pre-crRNA, 40 mM Tris-HCl pH 7.5, 60 mM NaCl unless stated otherwise. Typical reactions were incubated at 37°C for 30 minutes and quenched with 20 μL 2 loading and 100 °C for 5 minutes to stop the reactions. The products were separated by electrophoresis over 14% polyacrylamide gels containing 7 M urea and stained with Gel-Red, visualized by fluorescence imaging (pre-run at 600 V for 1 hour, sample run at 250 V for 30 minutes).

### MST binding assay

All microscale thermophoresis measurements (MST) were performed on a NanoTemper Monolith NT.115 instrument (NanoTemper Technologies, Munich, Germany). using the Standard Treated Capillaries K002 of the supplier. Each of the 16 solutions of one titration series was filled into a capillary, which were measured successively to create the respective data points in the experiment. General settings were applied for all MST experiments as follows: manual temperature control: 22°C, LED laser: RED. MST power is set at medium and excitation power is set at 20%.

For MST assay, all the proteins were exchanged into the MST buffer (20 mM HEPES pH7.5, 500 mM NaCl, 5% glycerol). The TPR-CHAT and its mutants were fluorescence-labeled using the Protein Labeling Kit RED-NHS 2nd Generation (NanoTemper Technologies) at 10 μM. 50 nM fluorescence-labeled TPR-CHAT and its mutants were added in a 1:1 ratio to a 1:2 dilution series with a final concentration of 2 μM down to 0.122 nM for gRAMP-crRNA complex. Each experiment was conducted at least 3 times and the similar result was obtained each time. Each protein *K_D_* value was obtained with a signal-to-noise ratio higher than 10. Datasets were processed with the MO (Monolith). Affinity Analysis v2.3 software. The analysis of dose-response curves was carried out with OriginPro 2018.

### Cryo-EM sample preparation

For cryo-EM sample preparation, the gRAMP-crRNA and Craspase complex were purified as described above. For the gRAMP-crRNA-TR and Craspase-TR complex, the purified gRAMP-crRNA or Craspase complex was incubated with TR with a molar ratio of 1:5 for 30 min. For the Craspase-NTR/CTR complex, the purified Craspase complex (gRAMP^D698A/D547A^) was incubated with NTR/CTR with a molar ratio of 1:5 overnight on the ice, then purified by size-exclusion chromatography to remove the excess RNA. All six samples were concentrated to 1.8 mg/mL for cryo-EM analysis. Next, 4 μL of protein samples were immediately applied to discharged 300-mesh Au R1.2/1.3 grids (Quantifoil, Micro Tools GmbH, Germany). Grids were blotted for 4 s and plunged into liquid ethane using an FEI Mark IV Vitrobot operated at 8°C and 100% humidity. The grids were stored in liquid nitrogen before data collection.

### Data collection and Image processing

The datasets of the gRAMP-crRNA, gRAMP-crRNA-TR, Craspase and Craspase-TR complex were collected on a Titan Krios microscope operated at a voltage of 300 kV by a Gatan K3 Summit direct electron detector with SerialEM software (36) at a nominal magnification of 105,000 × with a slit width of 20 eV. The pixel size was 0.855 Å/pixel. The defocus range were set from −1.7 μm to −2.5 μm. The electron exposure on the detector was about 40 e^−^/Å^2^. Each movie stack contains 40 frames. For the dataset of gRAMP-crRNA complex, 5,031 movies were collected. The beam-induced motion was corrected by MotionCor2 (37). The defocus values were estimated by CTFFIND 4.1 (38). A total of 3,081,589 particles were auto-picked on micrographs with dose-weighting using RELION 3.1 (39). Three rounds of 2D classification and two rounds of 3D classification was done with K = 5 classes and regularization parameter T = 20. An initial model was then generated by cryoSPARC v3 (40). 183,495 good particles were then subjected to per-particle CTF refinement (per-particle defocus and per-micrograph astigmatism estimation, as well as beam tilt estimation), followed by Bayesian particle polishing. 3D auto-refinement against the shiny particles were subjected to 3D classification without alignment with a generous mask with C1 symmetry, which was further improved to 3.01 Å by an additional round of CTF refinement (estimating both defocus, astigmatism and magnification anisotropy) and Bayesian particle polishing. A similar strategy was performed for the other three datasets. For the dataset of gRAMP-crRNA-TR complex, 4,727 movie stacks were acquired. 2,726,228 particles were auto-picked and 274,680 particles were yielded a map at 2.89 Å resolution. For the dataset of Craspase complex, 1,944 CTF-corrected cryo-EM images were manually selected. 1,141,519 particles were auto-picked and 229,997 good particles were improved a map at 2.88 Å resolution. For the dataset of Craspase-TR complex, 1,150,450 particles were auto-picked on 3,262 cryo-EM images and 143,181 particles were refined and postprocessed at 2.97 Å resolution.

The datasets of Craspase-NTR/CTR complex were collected on a Titan Krios microscope operated at a voltage of 300 kV by a Gatan K3 Summit direct electron detector with SerialEM software at a nominal magnification of 29,000 × with a pixel size of 0.97 Å and the defocus range was from −1.3 μm to −1.7 μm. The electron exposure on the detector was about 50 e^−^/Å^2^. Each movie stack contains 32 frames. Two datasets including 3,333 and 3,166 movies. Approximately two million particles were picked by blob-picker in RELION 3.1. The cryo-EM data processing was similar to the gRAMP-crRNA complex. Finally, 40,626 and 49,880 particles were subjected to 3D refinement and resolved to at 3.08 Å and 3.27 Å resolution. All reported resolutions are based on the gold-standard FSC=0.143 criteria, and the final FSC curves were corrected for the effect of a soft mask by using high-resolution noise substitution. The final density maps were sharpened by B-factors calculated with the RELION post-processing program. The final maps for model building and figure presentation were performed using DeepEMhancer (41). Further information for all samples is provided in Supplementary Table 1. Local resolution map was calculated using ResMap (42).

### Model building

Atomic models of the gRAMP and TPR-CHAT were predicted and modeled using AlphaFold2(30). We docked the corresponding models into our map by using UCSF chimera(43) and manually adjusted and re-built by COOT (44). The stereochemical quality of each model was assessed using MolProbity (45). All the figures were created in PyMOL (www.pymol.org), COOT and UCSF Chimera.

### Bacterial Growth Assays

Non-induced overnight cultures of bacteria (*E. coli* with pACYCDuet, pET28a and pETDuet, encoding gRAMP-RNA, CHAT, and Csx30-Csx31-RpoE, respectively) or negative control (*E. coli* with corresponding empty vectors) were diluted to a final OD_600 nm_ of 0.4 in LB medium supplemented with ampicillin, kanamycin, and chloramphenicol. 180 μL of the culture were then transferred into wells in a 100-well plate containing 18 μL of phage lysate (or 18 μL of SM buffer for uninfected control) for a final MOI of 4 and 0.1 for phage lambda and 2 μL of 10 mM IPTG (to a final IPTG concentration of 0.1 mM). Infections were performed in triplicates from overnight cultures prepared from separate colonies. Plates were incubated at 37°C with shaking in a Bioscreen C and an OD_600 nm_ measurement was taken every 15 min. Synthetic pre-crRNA contains spacers which are directed to the transcribed strand of early gene transcripts of the dsDNA phage λ. Direct repeat sequence was found in native CRISPR arrays from the *Candidatus* “Scalindua brodae” genome (GenBank: JRYO01000185.1).

Synthetic sequence of pre-crRNA (repeat marked in red): GTTATGAAACAAGAGAAGGACTTAATGTCACGGTACAGCTGCTCTTGTGTTAATGGTTTCTTTTTTGTGCTCATGTTATGAAACAAGAGAAGGACTTAATGTCACGGTACTTGTAGTCCTGAACGAAAACCCCCCGCGATTGGCACATGTTATGAAACAAGAGAAGGACTTAATGTCACGGTACCGCATTGCATAATCTTTCAGGGTTATGCGTTGTTCCATGTTATGAAACAAGAGAAGGACTTAATGTCACGGTACATTCGTAGAGCCTCGTTGCGTTTGTTTGCACGAACCATGTTATGAAACAAGAGAAGGACTTAATGTCACGGTACCCTCTGCCGAAGTTGAGTATTTTTGCTGTATTTGTCATGTTATGAAACAAGAGAAGGACTTAATGTCACGGTACCGGTCAAAGTTAACCATCTGTGCGGCGATGTTTTTCATGTTATGAAACAAGAGAAGGACTTAATGTCACGGTAC

For the experiments about bacterial cells expressing the target RNA, related plasmids harboring genes or their mutants were transformed into *E.coli* strain BL21(DE3). The cells were first grown to OD_600 nm_ of 0.6. Then 10-fold serial dilutions in LB were performed for each of the samples and 2.5 μL drops were put on agar plates containing 30 μg/mL chloramphenicol, 80 μg/mL ampicillin, 50 μg/mL kanamycin with or without 0.05 mM IPTG. Plates were incubated overnight at 37°C.

### Plaque assays

For the plaque assay, the related plasmids or its mutants were transformed into *E. coli* strain BL21(DE3), bacteria containing defense system and control bacteria with no system were grown overnight at 37°C. For cells that contained inducible constructs, added 0.2 mM IPTG induced 3 h at 37°C. And the inducers were added to the agar before plates were poured. Tittering was performed using 100 μL of E. coli cells grown to OD_600_ of 0.6 which induced 3h at 37°C. These cells were mixed with 100 μL of λ phage or 10-fold dilutions of each phage, incubated for 30 minutes and shaken every 10 min. 5 mL of molten 0.8% LB/agarose (pre-heated to 65°C) was added to the cell slurry and the soft agar was poured over the top of an LB-agar plate. Plates were incubated at 37°C for 12 h and then plaques were counted. Plaque forming units (PFUs) were determined by counting the derived plaques after overnight incubation and lysate titer was determined by calculating PFUs per mL. All experiments include three individual biological repeats with each including technical replicates in triplicate.

## Data and Materials Availability

The atomic coordinates for the Cryo-EM structures of the gRAMP-crRNA (PDB: 7XSO), gRAMP-crRNA-TR (PDB: 7XSP), Craspase (PDB: 7XSQ), Craspase-TR (PDB: 7XSR), Craspase-NTR (PDB: 7XT4) and Craspase-CTR (PDB: 7XSS) have been deposited in the Protein Data Bank (www.rcsb.org). The cryoEM density maps reported in this study, gRAMP-crRNA (EMD-33429), gRAMP-crRNA-TR (EMD-33430), Craspase (EMD-33431), Craspase-TR (EMD-33432), Craspase-NTR (EMD-33439) Craspase-CTR (EMD-33433) have been deposited in the EM Data Bank.

