## Supplementary files for "Structural and functional insights into the type III-E CRISPR-Cas immunity"

### Table S1. Cryo-EM data collection, refinement and validation statistics

| | gRAMP-crRNA | gRAMP-crRNA-TR | Craspase | Craspase-TR | Craspase-NTR | Craspase-CTR |
|---|---|---|---|---|---|---|
| **Data collection and processing** | | | | | | |
| EMDB code | EMD-33429 | EMD-33430 | EMD-33431 | EMD-33432 | EMD-33439 | EMD-33433 |
| PDB code | 7XSO | 7XSP | 7XSQ | 7XSR | 7XT4 | 7XSS |
| Electron microscopy | Titan Krios | | | | | |
| Camera | Gatan K3 Summit | | | | | |
| Magnification | 105,000× | | | | 29000× | |
| Voltage | 300 kV | | | | | |
| Defocus range (μm) | -1.3 to -2.3 | | | | | |
| Electron exposure (e-/Å$^2$) | 50 | | | | | |
| Pixel size (Å) | 0.855 | | | | 0.97 | |
| Exposure rate (e$^-$/A$^2$/sec) | 20 | | | | | |
| Number of frames per movie | 32 | | | | | |
| Energy filter slit width | 20 | | | | None | |
| Automation software | SerialEM | | | | | |
| Micrographs collected | 5,031 | 4,727 | 1,944 | 3,262 | 3,333 | 3,166 |
| Data processing software | RELION 3.1 & cryoSPARC 3.2 | | | | | |
| Symmetry imposed | *C*1 | *C*1 | *C*1 | *C*1 | *C*1 | *C*1 |
| Total extracted particles | 3,081,589 | 2,726,228 | 1,141,519 | 1,150,450 | 2,116,340 | 1,672,416 |
| Total refined particles | 183,495 | 274,680 | 229,997 | 143,181 | 40,262 | 49,880 |
| Final particles | 183,495 | 274,680 | 229,997 | 143,181 | 40,262 | 49,880 |
| Map resolution (FSC=0.143/Å) | 3.01 | 2.89 | 2.88 | 2.97 | 3.08 | 3.27 |
| Local resolution range (Å) | 2.0-4.0 | | | | | |
| **Refinement** | | | | | | |
| Initial Model (PDB code) | AlphaFold2 | | | | | |
| Refinement Package | Phenix 1.19 (real space refinement) | | | | | |
| Model resolution (FSC=0.5/Å) | 3.07 | 3.00 | 3.03 | 3.09 | 3.16 | 3.38 |
| Map sharpening B factor (Å$^2$) | -20 | -20 | -20 | -20 | -20 | -20 |
| Map CC | 0.82 | 0.81 | 0.81 | 0.80 | 0.81 | 0.80 |
| Model composition | | | | | | |
| Non-hydrogen atoms | 10,968 | 11,317 | 16,571 | 16,907 | 16,850 | 16,847 |
| Protein residues | 1,271 | 1,271 | 1,967 | 1,963 | 1,939 | 1,942 |
| Nucleotides | 35 | 50 | 34 | 49 | 55 | 54 |
| Ligands | 4 | 4 | 4 | 4 | 4 | 4 |
| *B* factors (Å$^2$) | | | | | | |
| Protein | 52.16 | 49.14 | 48.74 | 41.86 | 48.40 | 56.91 |
| Nucleotides | 46.50 | 53.78 | 42.82 | 33.68 | 42.77 | 54.13 |
| Ligands | 78.42 | 68.29 | 69.94 | 55.01 | 61.95 | 112.01 |
| R.m.s. deviations | | | | | | |
| Bond lengths (Å) | 0.009 | 0.007 | 0.009 | 0.007 | 0.005 | 0.003 |
| Bond angles (°) | 1.025 | 1.122 | 0.918 | 0.969 | 0.929 | 0.613 |
| **Validation** | | | | | | |

| | | | | | | |
|---|---|---|---|---|---|---|
| MolProbity score | 1.99 | 1.94 | 1.79 | 1.96 | 1.86 | 1.85 |
| EMRinger score | 3.04 | 1.81 | 3.06 | 2.87 | 3.42 | 2.22 |
| Clashscore | 3.64 | 3.50 | 2.51 | 3.23 | 3.76 | 8.40 |
| Poor rotamers (%) | 3.54 | 3.42 | 2.80 | 3.94 | 2.94 | 0.59 |
| C-beta deviations | 0.17 | 0 | 0.05 | 0 | 0.06 | 0 |
| CaBLAM outliers | 4.49 | 3.78 | 3.56 | 3.40 | 2.63 | 3.36 |
| Ramachandran plot (%) | | | | | | |
| Favored | 93.73 | 94.19 | 93.81 | 94.21 | 95.10 | 94.18 |
| Allowed | 6.20 | 5.65 | 6.14 | 5.69 | 4.69 | 5.72 |
| Outliers | 0.07 | 0.16 | 0.05 | 0.10 | 0.21 | 0.10 |

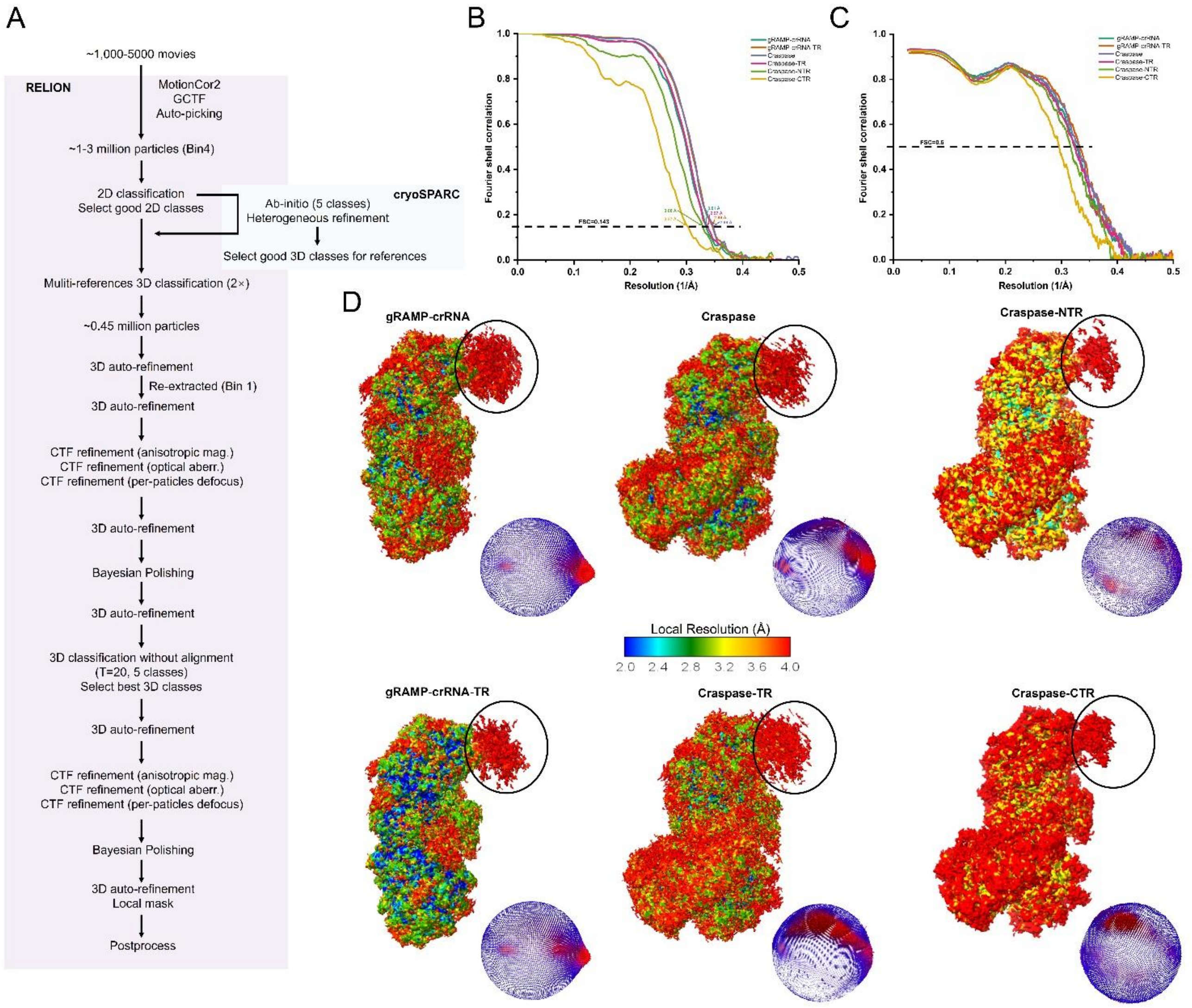


**Fig. S1 Single particle cryo-EM analysis of the samples in this study**

(A) A typical cryo-EM data processing workflow.

(B-C) Representative gold-standard Fourier Shell Correlation (FSC=0.143, left) and map vs model Fourier shell correlation curves (FSC=0.5, right) for each structure.

(D) Particle orientation distribution and local resolution-colored maps for each structure. Local resolution range was at 2.0-4.0 Å. The region corresponding to the insertion domain of gRAMP is marked with a circle.

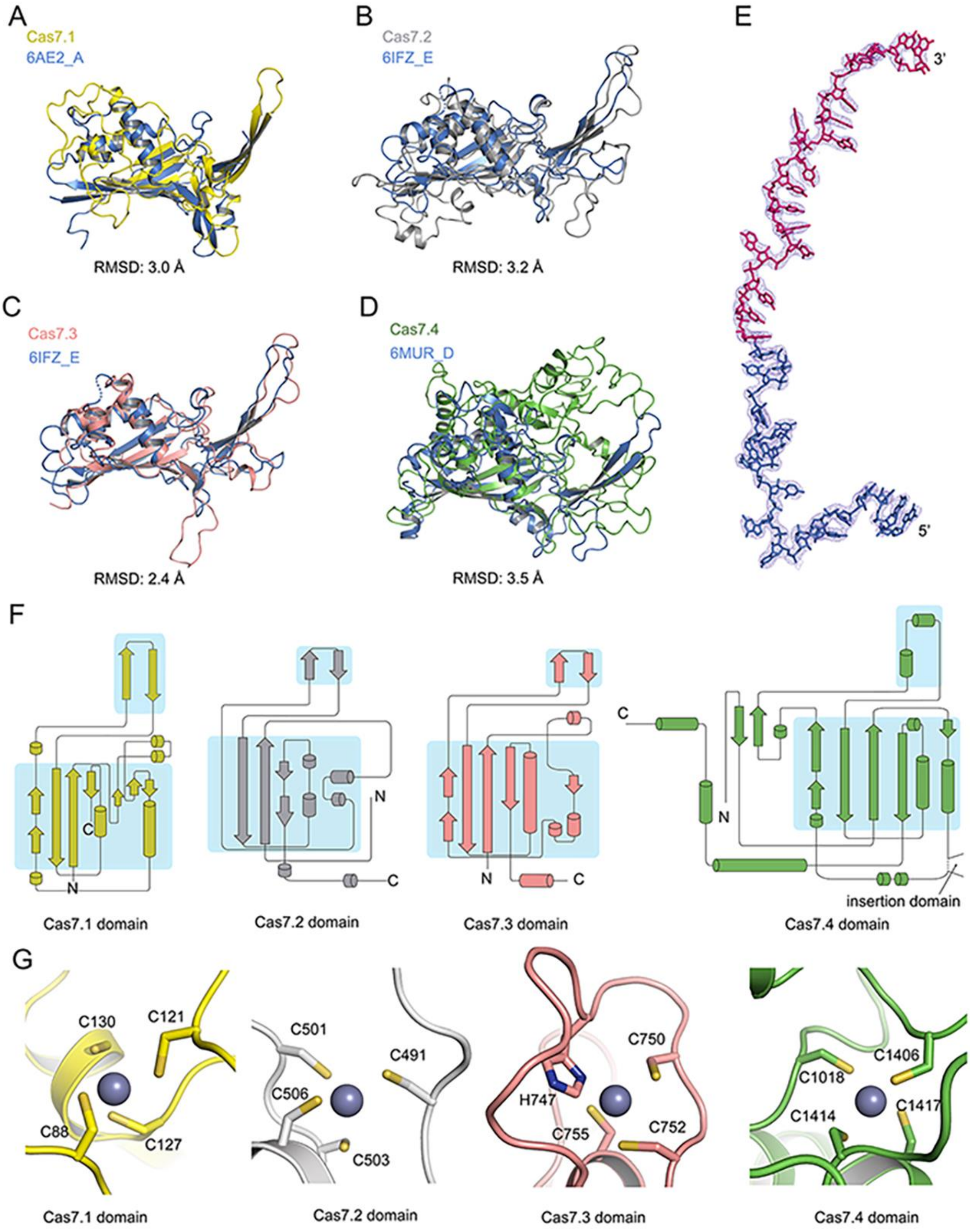


**Fig. S2 Structures of the Cas7 domains and crRNA**

(A-D) Structural superimpositions between Cas7 domains and their structural homologs. The PDB code and the chain number of the homologs are indicated and the RMSD values between the two superimposed structures are labeled below the structures.

(E) The structure of the crRNA within the apo gRAMP-crRNA with cryo-EM density shown in blue mesh.

(F) Topological diagrams of the Cas7.1-7.4 domains, in which the main body of Cas7 domains and the "thumb" motif are shaded.

(G) Binding of zinc ions in the zinc finger motif of Cas7.1-7.4 domains,

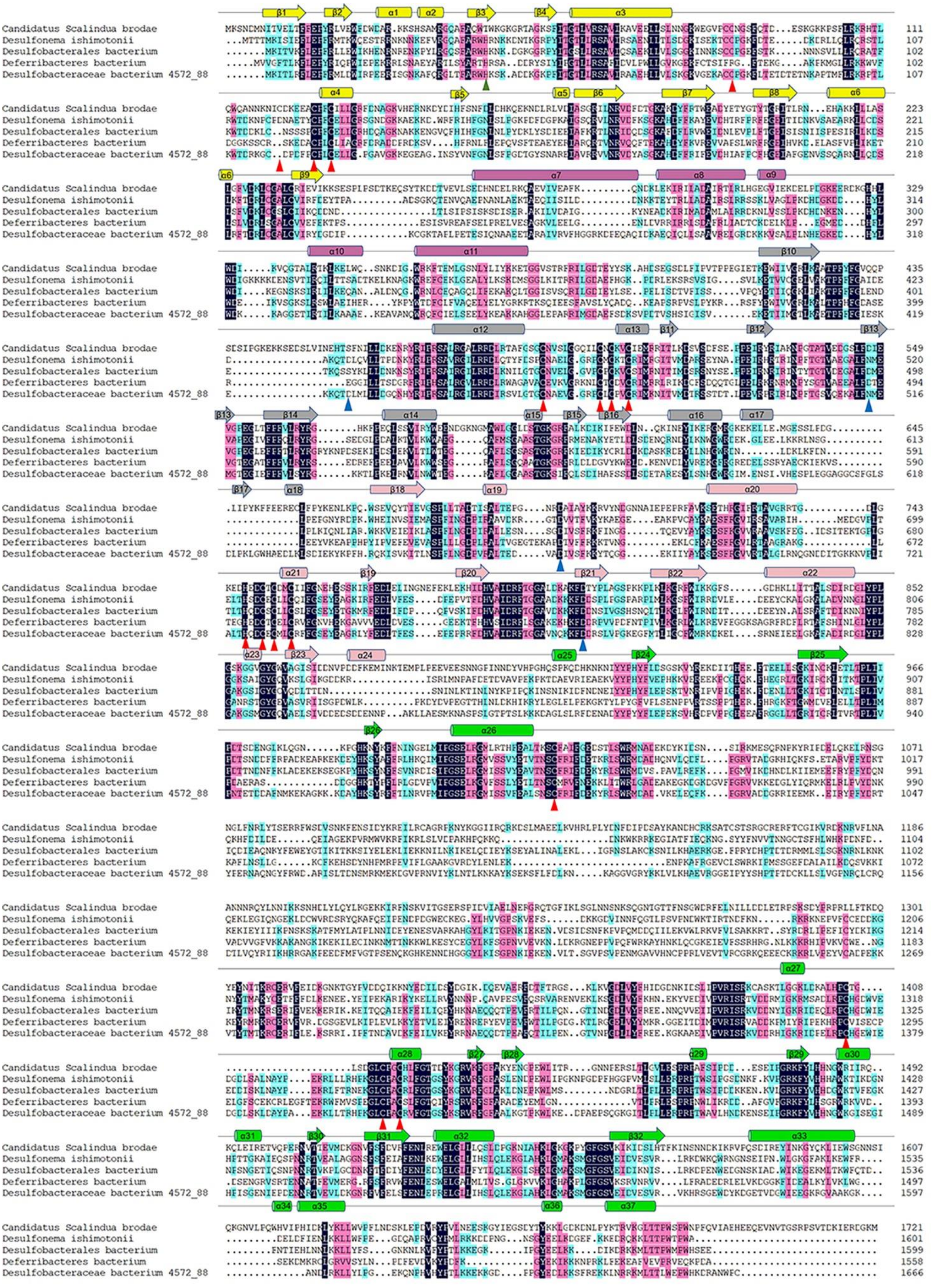

**Fig. S3 Sequence alignment of gRAMP homologs from different species**

Residues with 100 % identity, over 75 % identity and over 50 % identity are shaded in dark blue, pink and cyan, respectively. Secondary structural elements of Sb-gRAMP are shown above the sequences, colored as in Fig. 1A. gRAMP residues involved in zinc ion binding are marked with red triangles. Residues involved in target RNA cleavage and S457 are marked with blue triangles. The residue corresponding to H43 in Di-gRAMP is marked with a green triangle.

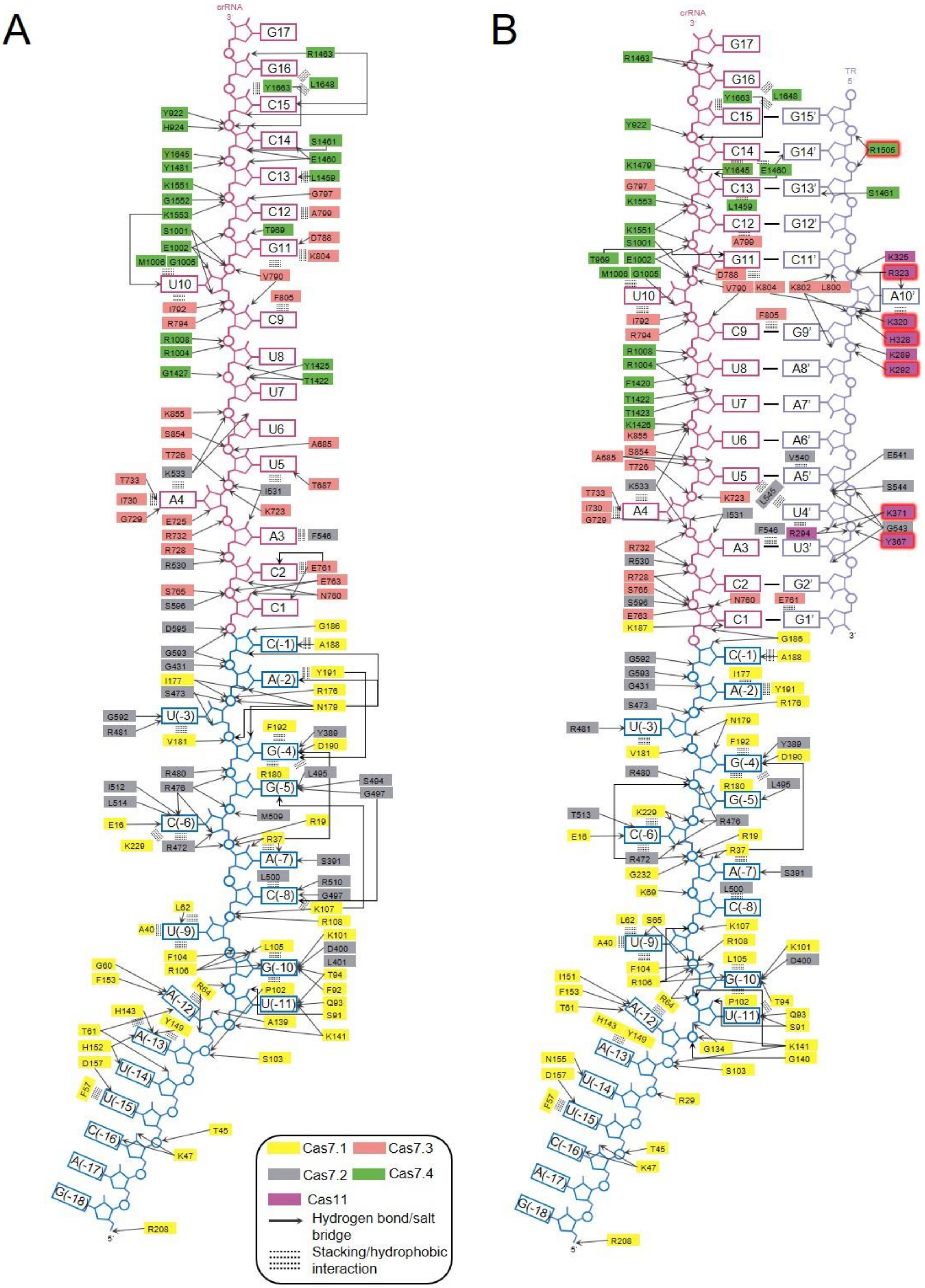


**Fig. S4 Nucleic-acid recognition in the gRAMP structures**

(A) Schematic view of the intermolecular interactions between gRAMP and crRNA in the apo gRAMP-crRNA structure.
(B) Schematic view of the intermolecular interactions between gRAMP and the crRNA- target RNA duplex in the gRAMP-crRNA-TR structure. Residues mutated in Fig. 2H are highlighted with red edges.

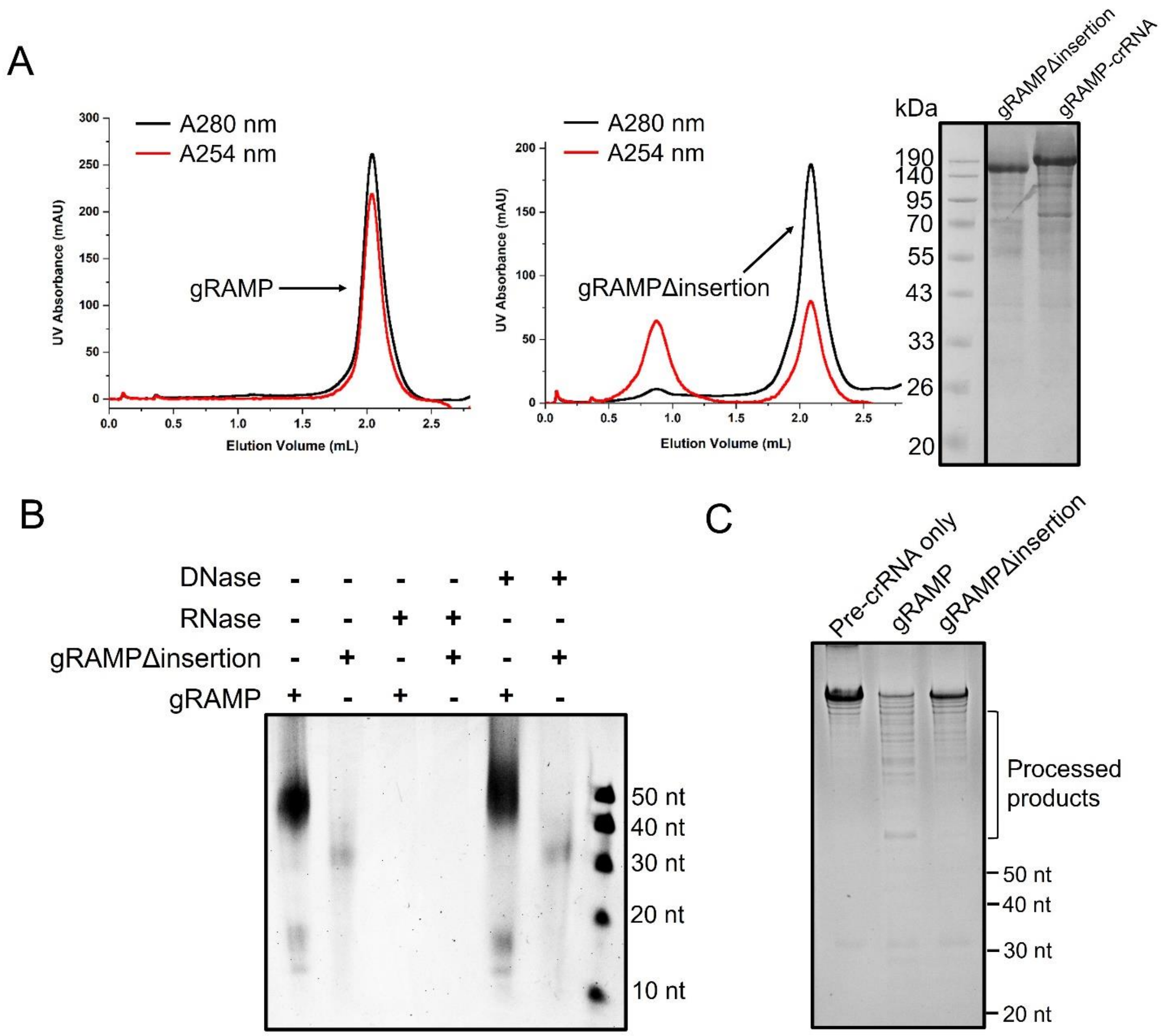


**Fig. S5 The insertion domain is involved in pre-crRNA processing**
(A) Gel filtration of gRAMP and gRAMPΔinsertion protein showing the absorptions at both 280 and 254 nm, respectively.
(B) Denaturing urea-PAGE gel showing the nucleic acids contained in the purified gRAMP and gRAMPΔinsertion co-expressed with CRISPR array. RNase, but not DNase can degrade the nucleic acids in purified gRAMP.
(C) Pre-crRNA processing using the transcribed CRISPR array with either purified gRAMP or gRAMPΔinsertion.

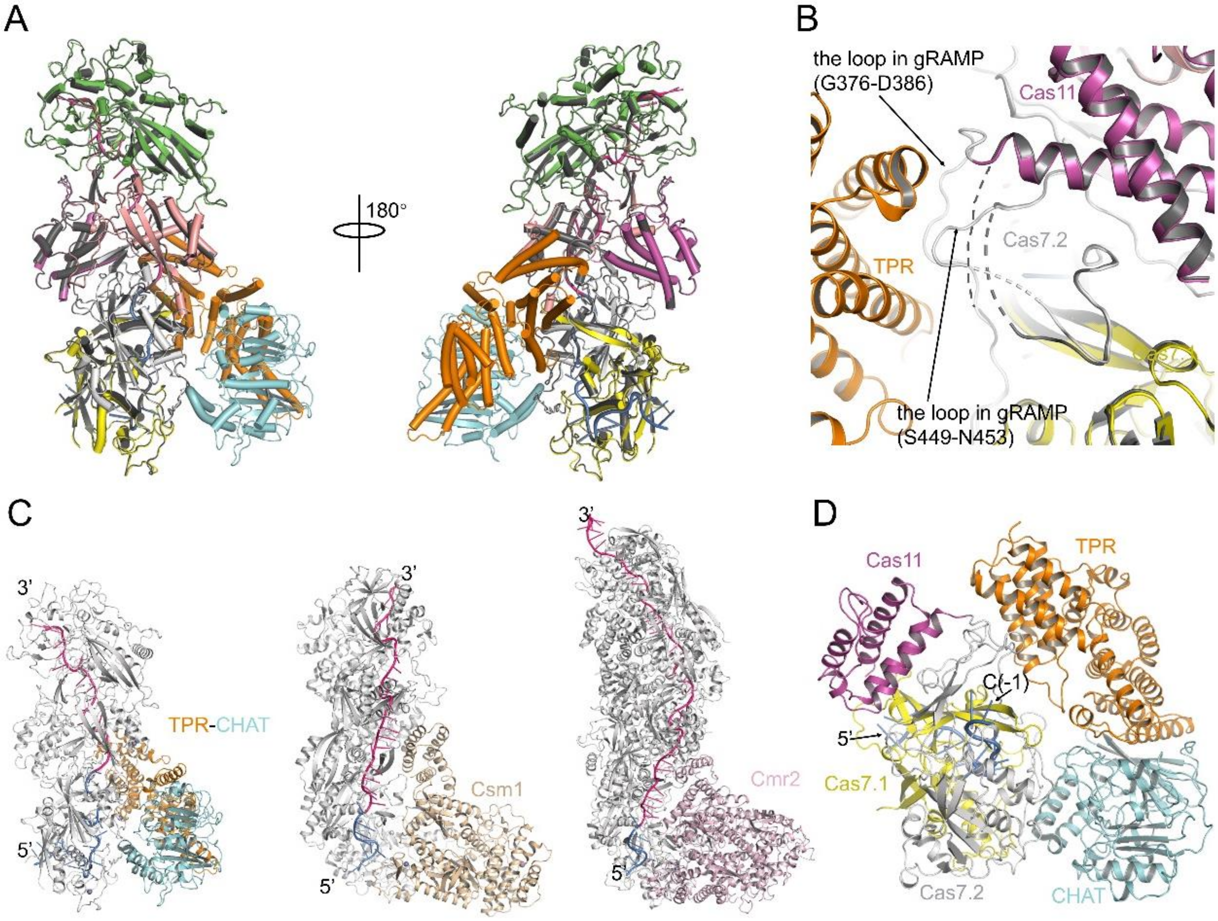


**Fig. S6 Cryo-EM structure of the apo Craspase structure**

(A) Structural superimposition between the gRAMP-crRNA (colored gray) and Craspsase (colored as in Figure 3A).

(B) The closed-up view of the superimposition in A showing that two loops in gRAMP become ordered in the structure of Craspase.

(C) Structural overviews of the type III-E, III-A and III-B CRISPR-Cas complexes showing that TPR-CHAT is situated at a similar position as Csm1 in type III-A and Cmr2 in type III-B complexes.

(D) The closed-up view of the crRNA position within the Craspase complex (colored as in Figure 3A).

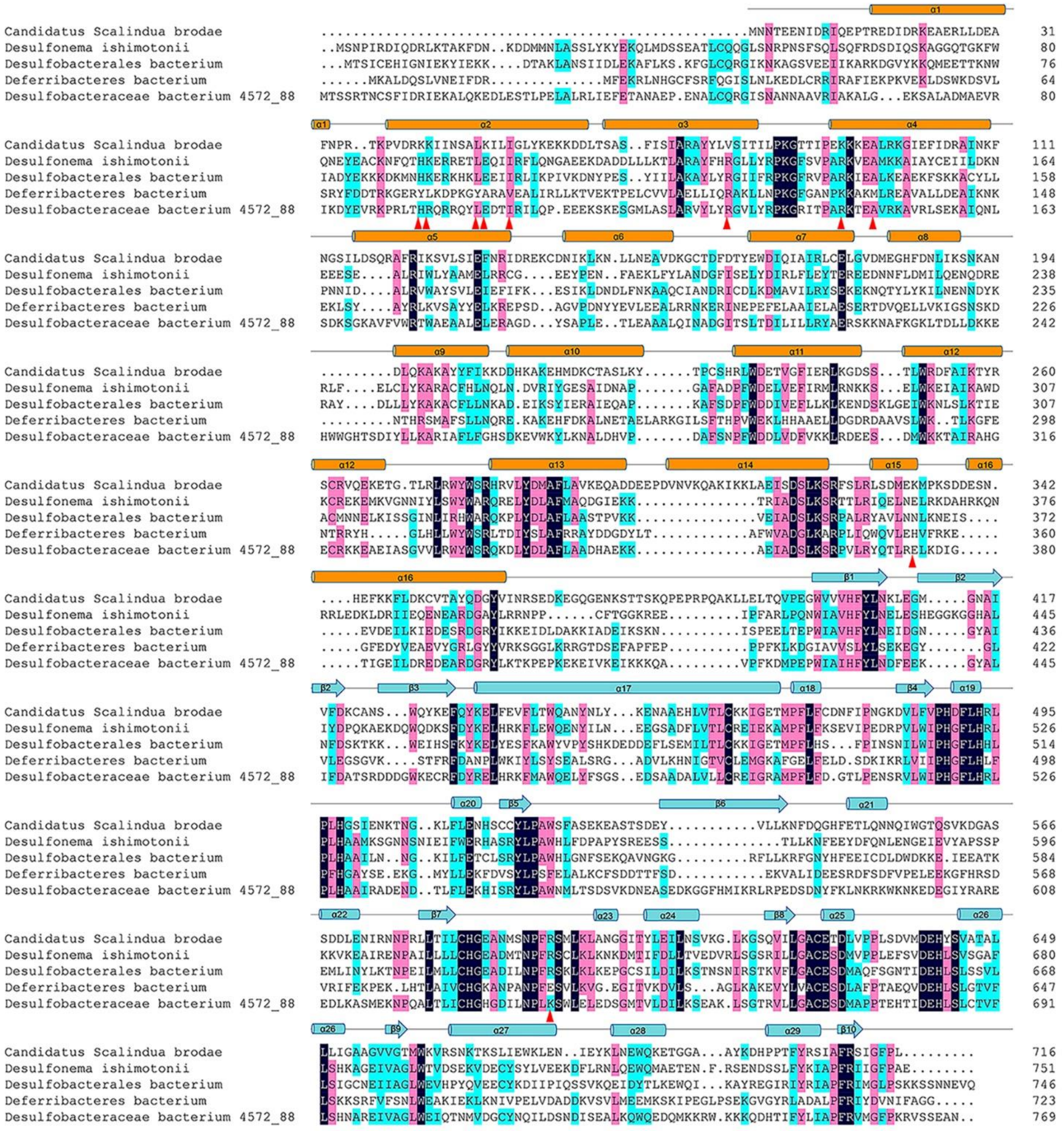

**Fig. S7 Sequence alignment of TPR-CHAT homologs from different species**

Residues with 100 % identity, over 75 % identity and over 50 % identity are shaded in dark blue, pink and cyan, respectively. Secondary structural elements of Sb-TPR-CHAT are shown above the sequences, colored according to the two domains, as in Figure 3A. Residues involved in gRAMP binding are marked with red triangles.

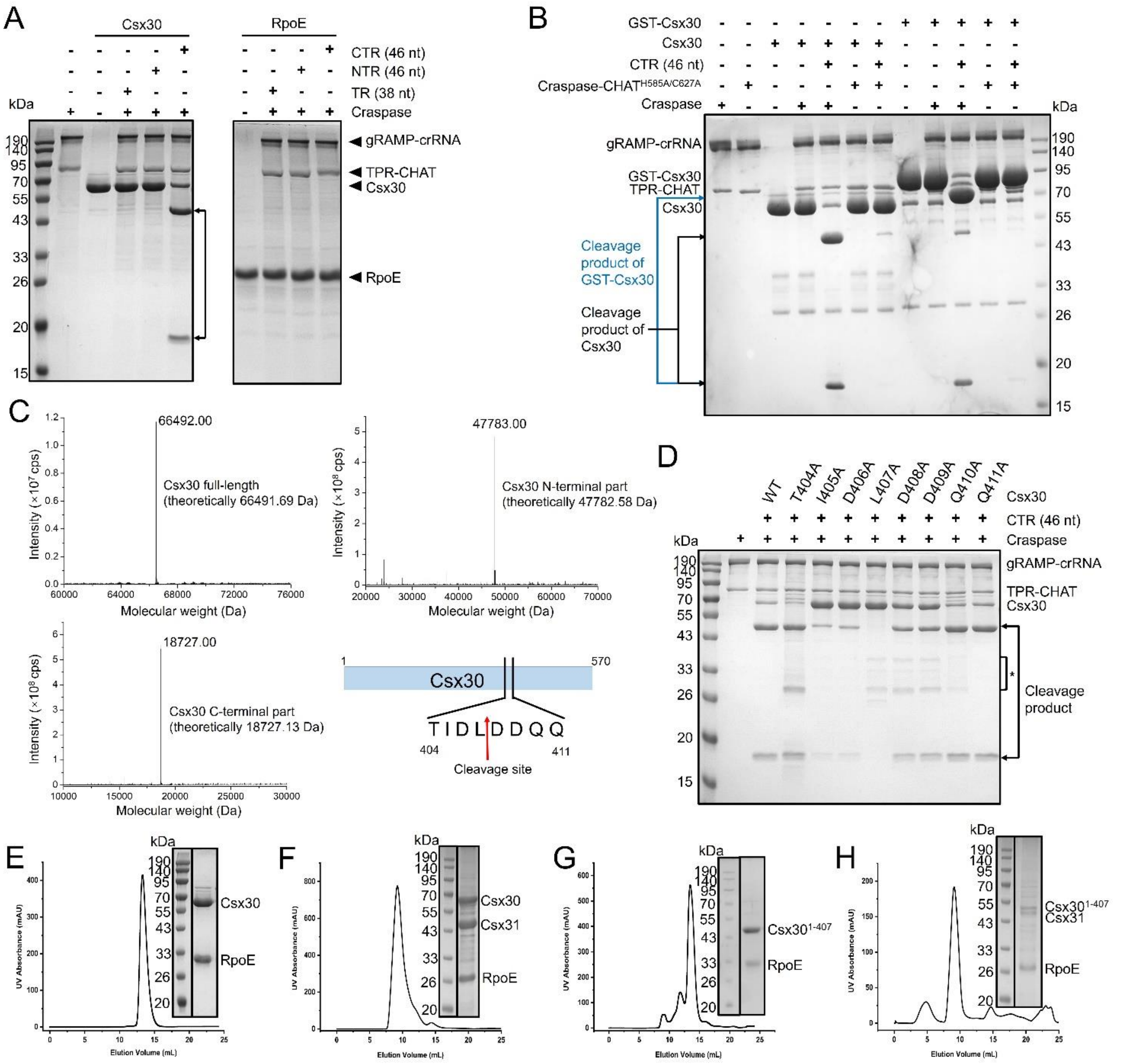


**Fig. S8 CTR binding activates the protease activity of Craspase towards Csx30**

(A) Protease activity assay of the Craspase bound by different target RNA molecules towards Csx30, Csx31 and RpoE.

(B) The target RNA dependent Csx30 cleavage by the TPR-CHAT in the Craspase complex. The TR/NTR/CTR RNA molecules, TPR-CHAT or its mutants, gRAMP-crRNA and Csx30 or N-terminally GST tagged Csx30 were added at 0.5, 0.3, 0.3 and 3 μM, respectively.

(C) LC-MS results of the purified full-length Csx30 and its two fragment products obtained from the protease cleavage. The intact molecular weight of the major protein samples are marked on the peak. The theoretical molecular weights of full-length Csx30, Csx30 (residues 1-407) and Csx30 (residues 408-570) are also indicated in the figures. A diagram showing the cleavage site is shown in the right bottom panel.

(D) The target RNA dependent Csx30 cleavage by the TPR-CHAT in the Craspase complex. The CTR RNA molecules, Craspase and Csx30 were added at 0.5, 0.3 and 1.5 μM, respectively.

(E-H) Gel filtration analysis of Csx30-RpoE (E), Csx30-Csx31-RpoE (F), $Csx30^{1-407}$-RpoE (G) and

Csx30$^{1-407}$-Csx31-RpoE complexes (H).

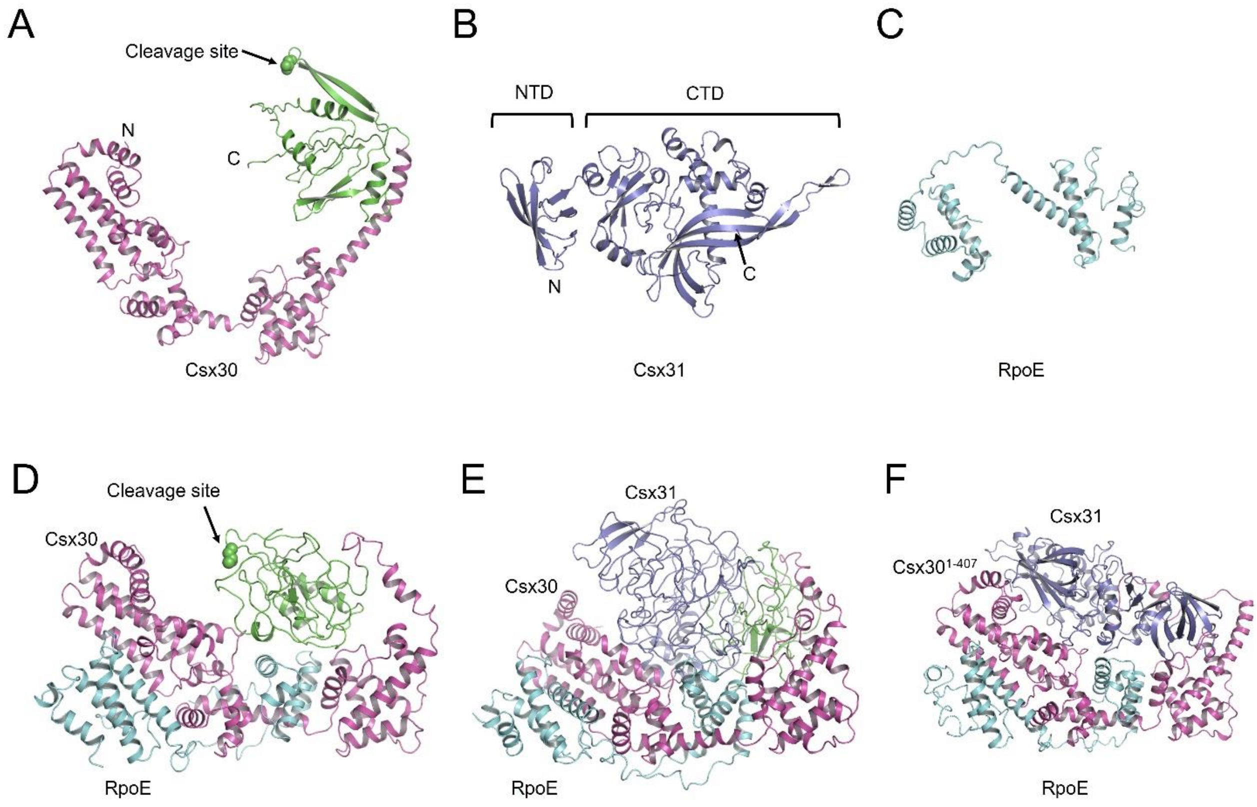


**Fig. S9 Structures of Csx30, Csx31, RpoE and their complexes predicted by AlphaFold2**

(A-C) Structures of Csx30 (A), Csx31 (B) and RpoE (C). The NTD and CTD of Csx30 is colored in magenta and green, respectively. The protease cleavage site is marked.

(D) Structure of the Csx30-RpoE complex. Csx30 and RpoE are colored as in A and C, respectively.

(E) Structure of the predicted Csx30-Csx31-RpoE complex.

(F) Structure of the predicted Csx30$^{1-407}$-Csx31-RpoE complex.

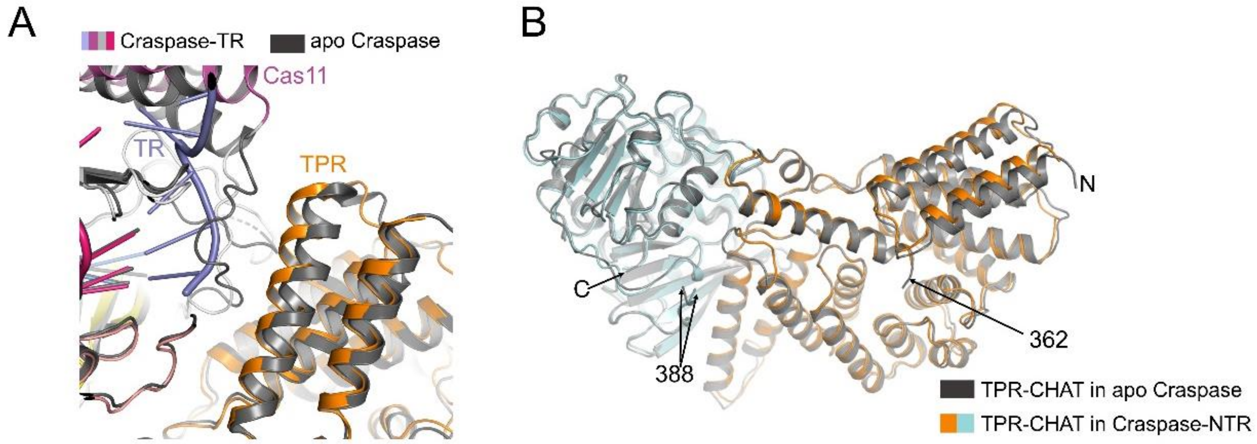


**Fig. S10 Conformational changes of TPR-CHAT caused by TR or NTR bound to the Craspase**

(A) Structural superimposition between Craspase (colored in dark gray) and Craspase-TR complex (colored as in Figure 5A).

(B) Structural superimposition of the TPR-CHAT molecule in the apo Craspase (colored in dark gray) and Craspase-NTR complexes (colored as in Figure 3A).

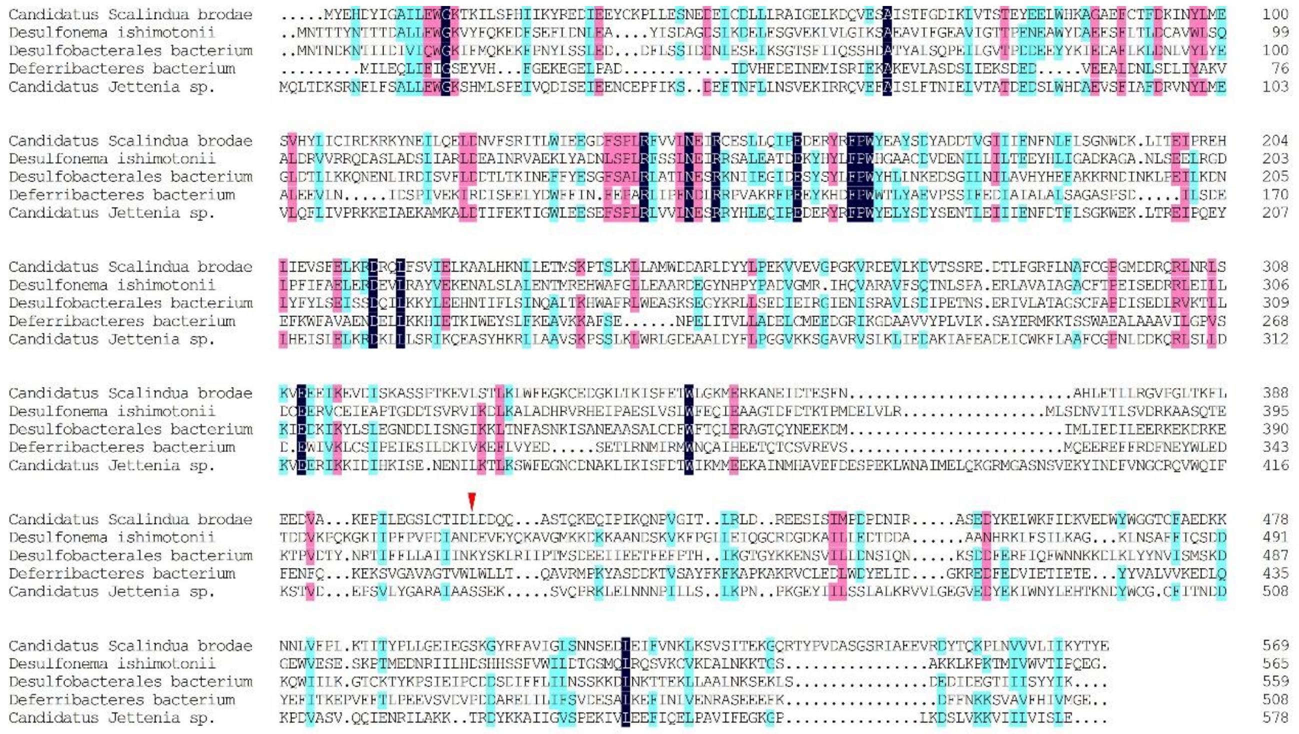

**Fig. S11 Sequence alignment of Csx30 homologs from different species**

Residues with 100% identity, over 75% identity and over 50% identity are shaded in dark blue, pink and cyan, respectively. The position of L407 in *Sb*-gRAMP is marked with a red triangle.
